# Targeted mutation of barley (1,3;1,4)-β-glucan synthases reveals complex relationships between the storage and cell wall polysaccharide content

**DOI:** 10.1101/2020.05.13.093146

**Authors:** Guillermo Garcia-Gimenez, Abdellah Barakate, Pauline Smith, Jennifer Stephens, Shi F. Khor, Monika S. Doblin, Pengfei Hao, Antony Bacic, Geoffrey B. Fincher, Rachel A. Burton, Robbie Waugh, Matthew R. Tucker, Kelly Houston

**Affiliations:** The James Hutton Institute, Invergowrie, Dundee, DD2 5DA, Scotland, UK; School of Agriculture and Wine, University of Adelaide, Waite Campus, Urrbrae, Adelaide, South Australia; La Trobe Institute for Agriculture and Food, School of Life Sciences, Department of Animal, Plant, and Soil Sciences, AgriBio Building, La Trobe University, Bundoora VIC 3086, Australia; Plant Sciences Division, College of Life Sciences. University of Dundee. Dundee, DD1 5EH, Scotland, UK

**Keywords:** barley, cell walls, gene editing, CRISPR/Cas9, (1,3;1,4)-β-glucan

## Abstract

Barley (*Hordeum vulgare* L) grain is comparatively rich in (1,3;1,4)-β-glucan, a source of fermentable dietary fibre that protects against various human health conditions. However, low grain (1,3;1,4)-β-glucan content is preferred for brewing and distilling. We took a reverse genetics approach, using CRISPR/Cas9 to generate mutations in members of the *Cellulose synthase-like* (*Csl*) gene superfamily that encode known (*HvCslF6* and *HvCslH1*) and putative (*HvCslF3* and *HvCslF9*) (1,3;1,4)-β-glucan synthases. Resultant mutations ranged from single amino acid (aa) substitutions to frameshift mutations causing premature stop codons, and led to specific differences in grain morphology, composition and (1,3;1,4)-β-glucan content. (1,3;1,4)-β-Glucan was absent in the grain of *cslf6* knock-out lines whereas *cslf9* knock-out lines had similar (1,3;1,4)-β-glucan content to WT. However, *cslf9* mutants showed changes in the abundance of other cell wall-related monosaccharides compared to WT. Thousand grain weight (TGW), grain length, width and surface area were altered in *cslf6* knock-outs and to a lesser extent TGW in *cslf9* knock-outs*. cslf3* and *cslh1* mutants had no effect on grain (1,3;1,4)-β-glucan content. Our data indicate that multiple members of the *CslF*/*H* family fulfil important functions during grain development but, with the exception of *HvCslF6*, do not impact the abundance of (1,3;1,4)-β-glucan in mature grain.

## Introduction

Cell walls are a distinctive structural feature of plant cells. They are composed of polysaccharide-rich layers that determine cell size and shape, and impact plant growth and development. The natural diversity of inter- and intra-species cell wall composition has an impact on the processing properties of plant-derived materials, both for industrial applications and human nutrition. For example, non-cellulosic polysaccharides are a key source of soluble dietary fibre that influence the nutritional quality and digestibility of plant-based foods (Burton and Fincher, 2012; Doblin *et al.*, 2010).

(1,3;1,4)-β-Glucan is a non-cellulosic polysaccharide found in the small grain cereals. Barley (*Hordeum vulgare* L) has higher grain (1,3;1,4)-β-glucan content (4–10% w/w) compared to other cereals such as wheat (*Triticum aestivum* L) (1% w/w) and rice (*Oryza sativa* L) (<0.06% w/w) (Burton and Fincher, 2012). A wide range of natural variation for grain (1,3;1,4)-β-glucan content (2.5–8% w/w) has been described in elite barley cultivars (Houston *et al.*, 2014; Izydorczyk *et al.*, 2000) and this influences their end-use properties. The brewing and distilling industries require barley cultivars with low grain (1,3;1,4)-β-glucan content for efficient malting and brewing. This is because (1,3;1,4)-β-glucan content has a direct impact on the viscosity of the mash and if too high, leads to filtration problems during brewing or to the formation of undesirable hazes in the final product (Gupta *et al.*, 2010; Wang *et al.*, 2004). Barley cultivars with high grain (1,3;1,4)-β-glucan content are preferred by the functional food sector based on its beneficial effect on human health (Cavallero *et al.*, 2002; Collins *et al.*, 2010; Keenan *et al.*, 2007). (1,3;1,4)-β-Glucan is not digested in the small intestine of humans, acting as fermentable dietary fibre that reduces the risk of diet-related conditions such as cardiovascular disease, type II diabetes and colorectal cancer (Brennan and Cleary, 2005; Wood, 2007; Cosola *et al.*, 2017; Ames *et al.*, 2018).

Members of three *Cellulose synthase-like* (*CslF/H/J*) subfamilies within the Glycosyltransferase (GT) 2 Family (Lombard *et al.*, 2014) synthesise (1,3;1,4)-β-glucan. However, the function of many members of each of these subfamilies are yet to be characterised (Little *et al.*, 2018). *HvCslF6*, *HvCslH1* and *HvCslJ* have been shown to be directly involved in (1,3;1,4)-β-glucan synthesis by either mutant knock-out/down of *CslF6* (Nemeth *et al.*, 2010; Vega-Sanchez *et al.*, 2012), transgenic overexpression of *HvCslF6* in barley grain via an endosperm-specific promoter (Burton *et al.*, 2011) or heterologous expression of *HvCslF6*, *HvCslH1* or *HvCslJ1* in either *Arabidopsis* or tobacco leaves (Little *et al.*, 2018). A genome-wide association study on grain (1,3;1,4)-β-glucan content (Houston *et al.*, 2014) confirmed that a QTL identified previously by Han *et al.* (1995) incorporates a cluster of genes including *HvCslF3*, *HvCslF4*, *HvCslF8*, *HvCslF10*, *HvCslF12* and *HvCslH1* on chromosome 2H, while a QTL on 1H co-locates with *HvCslF9* and *HvGlbI*, the latter being one of two (1,3;1,4)-β-glucan-specific endoglucanases. These QTL regions support the hypothesis that multiple *CslF* genes may act as mediators of (1,3;1,4)-β-glucan synthesis (Burton *et al.*, 2006) and contribute to natural variation in (1,3;1,4)-β-glucan content.

Although *HvCslF6* transcripts are the most abundant of all putative (1,3;1,4)-β-glucan synthase genes in developing grain and many vegetative tissues, *HvCslF9* is expressed during early grain development and in root tips (Burton *et al.*, 2008). *HvCslF3* is also highly expressed in root tips and coleoptiles. Due to the presence of (1,3;1,4)-β-glucan in the tissues where they are transcribed, both *HvCslF3* and *HvCslF9* are considered potentially capable of synthesising (1,3;1,4)-β-glucan (Aditya *et al.*, 2015; Burton *et al.*, 2008). In addition, the expression of *HvCslH1* in leaf tissue supports previous findings that this gene contributes to synthesis of (1,3;1,4)-β-glucan in this tissue (Burton *et al.*, 2008; Burton *et al.*, 2011; Doblin *et al.*, 2009). Despite distinct expression profiles, *in planta* function of the *HvCslF3, HvCslF9* and *HvCslH1* genes have yet to be confirmed in barley by loss-of-function mutants. This contrasts with the barley *CslF6* gene, where several *cslf6* alleles containing amino acid substitutions (*bgla, bglb, bglc*) have been described that exhibit reduced levels of (1,3;1,4)-β-glucan, susceptibility to chilling and alterations in grain morphology (Taketa *et al.*, 2012). Loss-of-function alleles for members of the *HvCslF/H* families would provide an opportunity to assess their role in (1,3;1,4)-β-glucan accumulation as well as plant growth and development.

Here we utilised a reverse genetics approach to provide insight into the contribution of each of four members of the *HvCslF/H* subfamilies to barley grain morphology and composition. We used CRISPR/Cas9-based gene editing technology to generate site-specific double strand breaks (DSB) leading to the introduction of targeted mutations via the non-homologous end joining (NHEJ) repair mechanism. We analysed plant and grain morphology/development, grain monosaccharide composition and the structure, distribution and content of (1,3;1,4)-β-glucan in the grain to provide insight into the function of each of the four genes. Our findings suggest that each of the genes contributes to the phenotype of barley grain, thereby providing opportunities to modify composition and morphology via *HvCslF/H* genes other than *HvCslF6*.

## Methods

### Plant material and *Agrobacterium*-mediated transformation

Barley cv. Golden Promise was used for embryo transformation via *Agrobacterium tumefaciens* following Bartlett *et al.* (2008) at the Functional Genomics (FUNGEN) facility, The James Hutton Institute (United Kingdom). Plants were grown in glasshouse conditions (16-h light/8-h dark photoperiod) until maturity in T_0_, T_1_ and T_2_ generations. At harvest, spikes were collected and manually threshed to obtain seed for subsequent generations.

### sgRNA design and construct assembly

sgRNA sequences were manually designed using the criteria stated on “Addgene CRISPR Guide” website (Addgene, 2018) to the 5’ end of the first exon for all four genes. This was to maximise the possibility of obtaining lines containing mutations that caused a knock-out of gene function induced by CRISPR/Cas9. To prevent off-target mutations, a BLAST search of sgRNA plus PAM sequences was performed on Barlex (Colmsee *et al.*, 2015) to determine sequence specificity to *HvCslF3*, *HvCslF6*, *HvCslF9* and *HvCslH1*, avoiding sgRNAs with potential matches to non-target genes and conserved protein motifs (Figure S4). Final sgRNA sequences were selected based on their target location and BLAST scores (Table S3 and S6). Each sgRNA was cloned into the pC95-gRNA entry vector downstream of the rice U6 small nuclear RNA (snRNA) (OsU6p) by Gibson Assembly. sgRNA for *HvCslF3*, *HvCslF6*, *HvCslF9* and *HvCslH1* were inserted into the pBract214m-*HvCas9*-HSPT expression vector which contains a barley codon optimised Cas9, and independently transformed into cv. Golden Promise (Method S1, Table S7, Figure S7).

### Screening of CRISPR/Cas9-induced mutations

Mutations were identified by a nested PCR method combined with InDel Detection by Amplicon Analysis (Yang *et al.*, 2015). Genomic DNA was isolated from a leaf disc (2 mm diameter) taken from individual barley seedlings using the Phire Plant Direct PCR Kit (Thermo Fisher Scientific, Waltham, USA). An aliquot of 0.5 μL crude plant extract was used as a template for *cas9* and InDel PCR detection in a reaction containing: 10 μL 2X Phire Plant PCR Buffer, 1 μL of each external forward and reverse primer at 10 mM (Table S8), 0.4 μL Phire Hot Start II DNA polymerase and 7.1 μL sdH_2_O in a total volume of 20 μL. External PCR amplicons, around 1.5 kb for each targeted gene, served as a DNA template (1 μL aliquot) for the nested (internal) PCR containing: 2.5 μL 10X Hot Start Taq buffer, 2.5 μL dNTPs, 0.12 μL Hot Start Taq DNA polymerase (Qiagen, Hilden, Germany) and 1 μL each primer: external forward and reverse at 1 mM and 10 mM, respectively and internal forward (6–FAM labelled) at 10 mM (Table S8) in a total volume of 25 μL. The internal forward (6–FAM) primer contains a 21-nucleotide tail complementary to the external forward primer. PCR conditions are described in Table S7. Fluorescent-labelled PCR amplicons were processed using an ABI3730 DNA Analyzer (Applied Biosystems Inc., Foster City, USA). The resulting chromatograms were visualised and analysed using GeneMapper^®^ (Applied Biosystems, v4.1) which allowed InDel identification based on size differences compared to cv. Golden Promise control. Mutations were confirmed by Sanger sequencing after reaction clean up and preparation as described in Houston *et al.* (2012). Resulting sequences were manually trimmed, aligned and analysed with Geneious V.9 (Kearse *et al.*, 2012) to identify sequence variations. Potential off-target effects were checked by PCR using primers described in Table S8, amplified using Hot Start Taq as described in Table S7. Only putative off-targets with zero mismatches in the PAM, i.e. *HvCslF3* and *HvCslH1*, where included as this motif is vital for Cas9 to recognise the sequence. DNA sequencing was carried out as described above.

### Phenotypic assessment of grain characteristics

Mature spikes were collected and hand threshed. A subset of ~30 bulked grains per genotype was used for phenotypic studies. All phenotypic data was collected from T2 plants (T3 grain). The number of grain, average grain area, length and width were measured with Marvin Seed Analyzer (GTA Sensorik GmbH, 2013). The grain were weighed to combine with the grain number estimate and derive thousand grain weight (TGW).

### Grain (1,3;1,4)-β-glucan quantification

For each genotype, five mature grains were milled and used to quantify (1,3;1,4)-β-glucan content using a modified version of the Mixed Linkage β-Glucan Assay Kit (K-BGLU, AACC Method 32-23.01, AOAC Method 995.16, Megazyme Int. (Wicklow, Ireland)) and based on the method described by McCleary & Codd, (1991) for the analysis of small flour samples (15 mg) described in Burton *et al.* (2011). Grain (1,3;1,4)-β-glucan content from two to four independent genotypes was averaged for each type of CRISPR/Cas9-induced mutation and calculated as % of dry weight (w/w). Each batch of grain (1,3;1,4)-β-glucan assays included a standardized barley flour control (4.1% w/w of (1,3;1,4)-β-glucan, from Megazyme kit) and duplicate technical replication.

### Immunolabelling of CRISPR/Cas9 gene-edited lines

Five grains per genotype were fixed in 4% (v/v) paraformaldehyde in PEM buffer solution [PEM Buffer = 0.1 M PIPES (pH 6.95), 1 mM EGTA, 1 mM MgSO4] for 24 hrs at 4°C. After fixation, samples were washed with ethanol (40% and 70%), transferred to a Leica TP1020 tissue processor and wax embedded in Paramat Extra, in pastille form (VWR International, Lutterworth, UK). Transverse grain sections (10 μm) were obtained using a RM2265 Rotary Microtome (Leica Biosystems, Vista, USA) and de-waxed prior to antibody labelling. For (1,3;1,4)-β-glucan detection, a 1:50 dilution of anti-mouse primary antibody, BG1, (Sigma-Aldrich, St. Louis, USA) was used. Goat anti-mouse secondary antibody conjugated to Alexa Fluor 488 (Sigma-Aldrich, St. Louis, USA) was used for fluorescence detection, diluted 1:100. Each batch of immunolabelling assays contained two cv. Golden Promise negative controls, no primary antibody with secondary and vice versa performed in parallel with the rest of the samples. Transverse grain sections were counterstained with 0.001% Calcofluor White (Sigma-Aldrich, St. Louis, USA). For starch staining, a 1:5 dilution of Lugol’s iodine solution (Sigma-Aldrich, St. Louis, USA) was applied to the grain sections. Lignin and differential polysaccharide staining were detected using a 0.02% toluidine blue O solution (Sigma-Aldrich, St. Louis, USA) as described in Methods S2.

### Imaging

Imaging of immunolabelled grain sections was performed on a Zeiss LSM 710 Confocal Laser Scanning Microscope (Carl Zeiss Microscopy GmbH, Jena, Germany) using a PL APO 20X /1.0 water dipping objective (Zeiss). Calcofluor was excited using 405 nm light from a blue diode laser and emission was collected between 420 and 460 nm. Alexa Fluor 488 dye (BG1) was excited using 488 nm laser light from an argon ion laser and emission was collected between 504 and 543 nm. Grain autofluorescence was detected by excitation using 594–633 nm laser light from a red diode laser and emission was collected between 631 and 730 nm. Cultivar Golden Promise with 5.00% w/w ± 0.03 of grain (1,3;1,4)-β-glucan content, was used as a positive control for (1,3;1,4)-β-glucan detection. Negative controls showed no fluorescent labelling when neither the primary antibody nor secondary were used (Figure S8). Bright field images of grain sections stained with Lugol’s iodine solution and toluidine blue O were collected using a Zeiss Axioskop 2 Plus microscope (Carl Zeiss Microscopy GmbH, Jena, Germany) equipped with a 5X lens (Zeiss). Subsequent to collection, images were processed with Zen Software v6.0., utilizing global adjustment tools only.

### DP3:DP4 analysis

The DP3:DP4 ratio was determined by High-Performance Anion-Exchange Chromatography (HPAEC) as described in Ermawar *et al.* (2015). Lichenase enzyme (Megazyme Int., Wicklow, Ireland) was used to release oligosaccharides for profiling by HPAEC. A no-enzyme treated control, a lichenase digest of barley flour, laminaribiose and laminaritriose were included as controls under the same chromatographic conditions as for CRISPR/Cas9-induced lines (Doblin *et al.*, 2009; Pettolino *et al.*, 2009).

### Non-cellulosic monosaccharide analysis

Non-cellulosic monosaccharide analysis was carried out as described in Hassan *et al.* (2017) using reversed phase HPLC separation coupled to diode array detection (Agilent Technologies). Dehulled grain was ground and resulting flour samples (20 mg) were prepared according to Pettolino *et al.* (2012). Acid hydrolysis of non-destarched alcohol insoluble residue (AIR) was undertaken by adding 1 M sulphuric acid, as described previously (Burton *et al.*, 2011), to the insoluble material (~15 mg). Between 3 and 12 replicates were analysed for each genotype. The moisture content of the ground flour was determined as outlined in AOAC 925.10 (AOAC International, 2005). All analyses were performed using quadruplicate technical replication.

## Results

### CRISPR/Cas9 induced mutations in members of the *HvCslF/H* gene family

We catalogued the mutations in each of the target genes over three generations (T_0_, T_1_ and T_2_). In the T_0_ plants all CRISPR/Cas9-induced mutations were in the heterozygous state. The frequency of mutations ranged from 35.3% for *HvCslF3*, to 9.3% for *HvCslF6*, 22% for *HvCslF9* and 4.7% for *HvCslH1* (Table S1). In putative T_0_ mutants, 83% of the plants carried short InDels (ranging from 1 bp deletions to 2 bp insertions; Table 1, Figure S1). An exception to this trend was identified for *HvCslF9* where a 39 bp deletion was detected in the heterozygous state in a single T_0_ plant.

**Table 1.** Collection of *cslf*/*h* mutants (T_2_) used for phenotypic characterisation in this study and their predicted protein effect. The mutation location was described with respect to the cDNA start. PAM represents Proto adjacent spacer motif, aa represents amino acid. Mutation locations are provided as nucleotides from the start codon followed by “ins” for an insertion or “del” for a deletion.

| Gene | InDel size | Allele | Allelic state | Type of mutation | Mutation location | Predicted protein effect |
| --- | --- | --- | --- | --- | --- | --- |
| *HvCslF3* | -2 bp | *cslf3-1* | Homozygous | Frameshift | 148_150del | Multiple aa changes |
|  | +3 bp | *cslf3-2* | Homozygous | In frame | 149_152ins | 1 aa insertion |
| *HvCslF6* | +1 bp | *cslf6-2* | Homozygous | Frameshift | 177_178ins | Premature stop codon (28 aa downstream PAM) |
|  | +1 bp | *cslf6-2/+* | Heterozygous | Frameshift | 177_178ins | Semi functional (Premature stop codon in one allele, 28 aa downstream PAM) |
| *HvCslF9* | -39 bp | *cslf9-1* | Homozygous | In frame | 117_156del | 13 aa deletion |
|  | -5 bp | *cslf9-2* | Homozygous | Frameshift | 126_131del | Premature stop codon (2 aa downstream PAM) |
|  | +1 bp | *cslf9-3* | Homozygous | Frameshift | 130_131ins | Premature stop codon (2 aa downstream PAM) |
| *HvCslH1* | -21 bp | *cslh1-1* | Homozygous | In frame | 25_46del | 7 aa deletion |
|  | +1 bp | *cslh1-2* | Homozygous | Frameshift | 48_49ins | Multiple aa changes |
|  | +13 bp | *cslh1-3* | Homozygous | Frameshift | 52_65ins | Multiple aa changes |

Homozygous lines were detected in the T_1_ (14 out of 58 for *HvCslF3*, 8 out of 169 for *HvCslF9* and 2 out of 58 for *HvCslH1*). These lines contained InDels of varying sizes, some of which had the potential to change the protein sequence. Screening for the presence of Cas9 by PCR indicated a high prevalence of transgene retention in *HvCslF6* and *HvCslH1* genotypes (92% and 97%, respectively) compared to *HvCslF3* (55%) and *HvCslF9* (56%) in T_1_ plants (Table S2). A range of T_1_ alleles were selected to maximise the possibilities of identifying potential phenotypic and allelic changes in the subsequent generation. A subset of 10 T_2_ alleles (Table 1, Figure 1; *cslf3*, *cslf6*, *cslf9* and *cslh1*) were used to characterise the effect of these mutations on grain morphology and composition.

**Figure 1.**
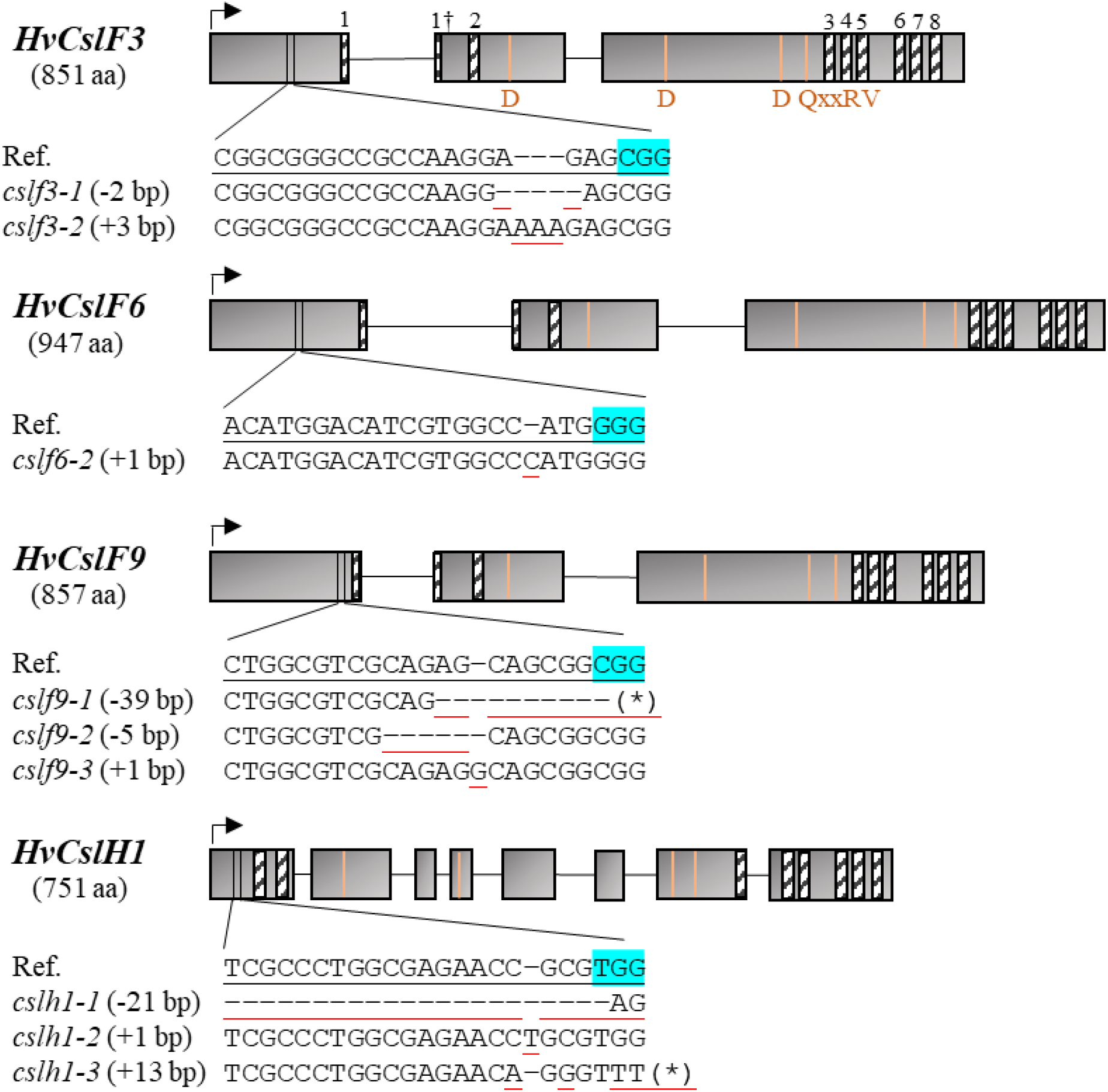
Collection of gene-edited *cslf/h* mutants used for phenotypic analysis of the grain. Exons are represented by grey boxes. Black arrows represent start codon. Black and white bars indicate the positions of eight predicted transmembrane domains (TM) across all genes (TM 1† expands through the first intron of *HvCslF* genes). Orange bars represent the D, D, D, QxxRW motifs characteristic of the GT_2_ Family. For each gene, the sgRNA reference sequence (Ref.) is underlined in black. InDels are underlined in red. PAM motif is highlighted in blue. Asterisks (*) represent 31 deleted nucleotides downstream the PAM motif in *HvCslF9* and a 13 bp insertion in *HvCslH1*.

### Mutations in *HvCslF/H* genes influence grain morphology

Using grain from a subset of 10 T_2_ lines of *cslf3 (2)*, *cslf6 (2)*, *cslf9 (3)* and *cslh1 (3)* we assessed the effect of each mutation on thousand grain weight (TGW), grain area, grain length and grain width and compared these to wild type (WT) grain of cv. Golden Promise. Cas9-free T_2_ lines were identified for *cslf3*, *cslf9* and *cslh1*, however, multiple Cas9 insertions were suspected in *cslf6-2* lines based on a preliminary segregation test for the selectable marker (Figure S2).

Grain from the *cslf6-2* mutants containing a premature stop codon (Table 1) in the first exon exhibited a significantly lower TGW (p = 0.002, Tukey’s test) compared to WT (Figure 2B). In addition, the grain was significantly longer (p = 0.006, Tukey’s test) compared to WT. In heterozygous mutants (*cslf6-2/+*), TGW was affected to a lesser extent, due to higher phenotypic variation for this trait. Grain were significantly longer (p = 0.002, Tukey’s test) than WT, whereas grain width was not significantly different, possibly leading to larger grains in terms of overall grain area. Significant differences in grain from *cslf6* homozygous and heterozygous plants compared to WT indicates there may be some dosage dependency of *HvCslF6* in sporophytic (maternal) tissues, similar to that reported previously for substitution alleles (Taketa *et al.*, 2008).

**Figure 2.**
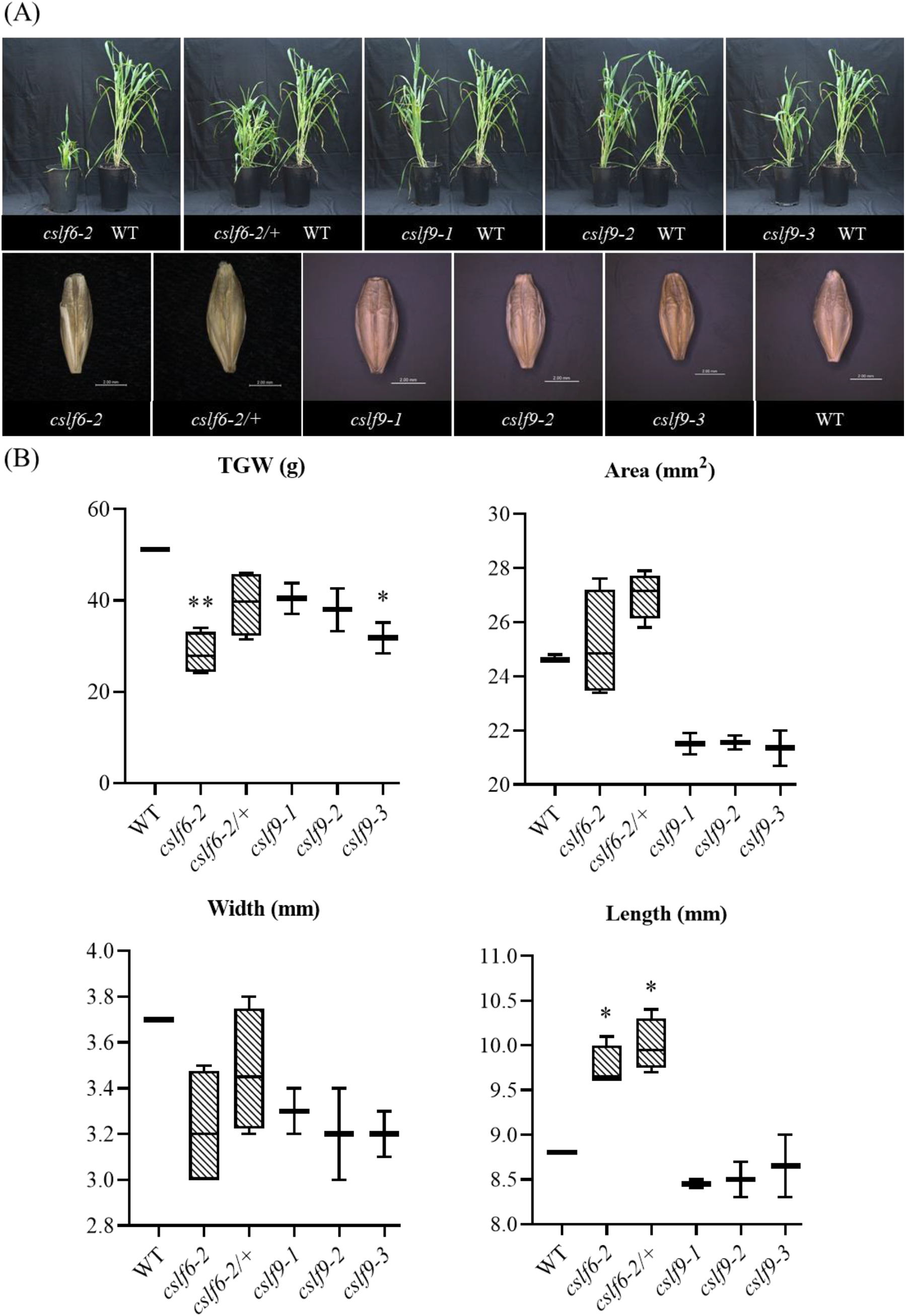
Whole plant and grain morphology of *cslf6* and *cslf9* alleles **(A)** Plant phenotype of *cslf6* (*cslf6-2* and *cslf6-2/+*) and *cslf9* (*cslf9-1*, *cslf9-2* and *cslf9-3*) gene-edited lines 10 weeks post germination. Each gene-edited line (left) was compared to cv. Golden Promise (WT; right). Below, mature grains harvested from the same *cslf6* and *cslf9* gene-edited lines. **(B)** Phenotypic assessment of grain characteristics: thousand grain weight (TGW), overall grain area, grain width and grain length for *cslf6* and *cslf9* lines. Error bars represent SD from two to four independent genotypes per mutation. Asterisks indicate statistically significant differences compared to the control (WT) at p < 0.05 (*) and p < 0.01 (**) using Tukey’s post-hoc test in GraphPad Prism v.8.4.2.

The *cslf9* homozygous mutant (*cslf9-3)* that carries a premature stop codon in the first exon (due to a 1 bp insertion) had significantly lower TGW compared with WT (p = 0.021, Tukey’s test; Figure 2B) whereas no significant differences were observed in *cslf9-1*, carrying a 39 bp in-frame deletion and *cslf9-2* carrying a 5 bp deletion leading to a premature stop codon also in the first exon (p =0.310; *cslf9-1* and p = 0.154; *cslf9-2*, Tukey’s test; Figure 2B). No statistically significant differences were identified for grain length (p = range: 0.1694–0.987, Tukey’s test, *cslf9* mutants vs WT) or overall area (p range = 0.087–0.116, Tukey’s test) compared to WT. However, grain from *cslf9* homozygous mutants had significantly smaller overall area compared to either *cslf6-2* (p = 0.038; *cslf9-1*, p = 0.041; *cslf9-2*, p = 0.030; *cslf9-3*, Tukey’s test) or *cslf6-2/+* mutants (p = 0.002; *cslf9-1*, p = 0.003; *cslf9-2*, p = 0.002; *cslf9-3*, Tukey’s test).

The *cslf3* mutants *cslf3-1* and *cslf3-2*, which carry a 2 bp deletion and a 3 bp insertion that cause a frame shift and the addition of a single amino acid respectively (Table 1), showed a significant reduction in TGW (p = <0.022 and 0.002, Tukey’s test) but to a lesser extent than the *cslf6* and *cslf9* mutant lines (Figure S4). *cslf3-2* had significantly narrower grains compared to WT (p = 0.008, Tukey’s test) although no significant differences were observed in grain area between *cslf3* mutants and WT grain (p = 0.261; *cslf3-1* and p = 0.157; *cslf3-2*, Tukey’s test; Figure S3). A lower TGW was observed for *cslh1* homozygous frameshift mutants (*cslh1-2* and *cslh1-3*) compared to WT (p = 0.012; *cslh1-2* and p = 0.044; *cslh1-3*, Tukey’s test; Figure S5). No differences were observed in relation to overall area (including grain width and length) for *cslh1* mutants compared to WT grain (p range = 0.243–0.648, Tukey’s test).

Based on BLAST searches, the sgRNA sequences for *HvCslF3* and *HvCslH1* have potential off-targets that could result in mutations being introduced into additional genes (Table S3). To assess if these off-target mutations were present in the lines characterised here we amplified the putative off-target amplicons in the same 10 lines (four *cslf3* lines and six *cslh1* lines) used for phenotypic characterisation of the grain (Table S4). For *HvCslF3* the sgRNA could potentially bind to *HORVU2Hr1G042350* (encoding a putative *Cellulose synthase-like D2* based on gene annotation), and for *HvCslH1* the potential off-target site was in *HORVU2Hr1G074940* (encoding a putative *Cellulose synthase-like B4)*, however, no mutations were observed in these genes. There were no putative off-targets for the sgRNAs designed to *HvCslF6* and *HvCslF9* because any potential matches identified contained mismatches within the PAM (Figure S6). Therefore, we conclude that for all the lines characterised that the phenotypes observed are due to mutations in the targeted genes.

### Impact of the induced mutations on plant development

The same lines used for phenotypic characterisation of grain development were analysed for changes in whole plant growth under glasshouse conditions. Due to the small number of seed available for some lines it was not feasible to carry out a full replicated study of gross plant morphology during development. However, several phenotypes were striking, and therefore worth describing here.

We observed a delay in development and a reduction in plant height in both *cslf6-2* (homozygous) and *cslf6-2/+* (heterozygous) mutants compared to WT (cv. Golden Promise). *cslf6-2* mutants had fewer tillers than WT after 10 weeks (Figure 2A). Five weeks later, spikes were developing in the cv. Golden Promise control whereas the *cslf6-2* mutant line was still in the vegetative phase. At maturity, only two to three spikes set seed in the *cslf6-2* mutants, limiting further grain phenotypic studies (Figure S3). The phenotypic appearance of *cslf6-2* homozygous grain was vastly different to other *csl* mutants (~50% reduction in TGW; Figure 2 and 3C). No major whole plant phenotypic changes were detected in *cslf9* mutants compared to WT (Figure 2A). A similar plant phenotype to WT was observed in *cslf3* and *cslh1* gene-edited lines (Figure S4A and S5A).

**Figure 3.**
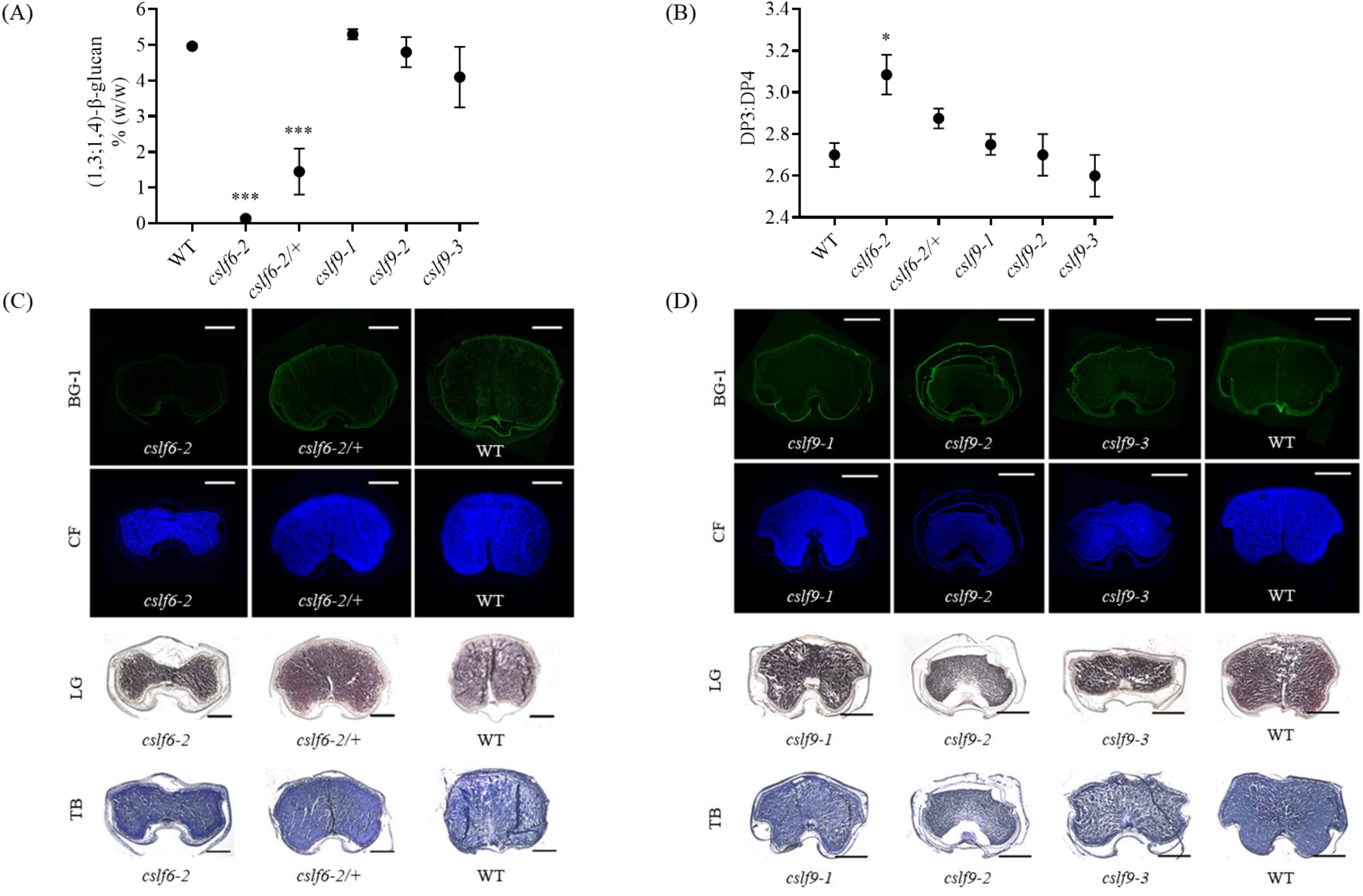
**(A)** Quantification of (1,3;1,4)-β-glucan concentration in mature grain of *cslf6* and *cslf9* mutants. **(B)** DP3:DP4 ratio of (1,3;1,4)-β-glucan in mature grain of the same lines. Error bars represent SE from two to four independent genotypes per mutation, except for a single *cslf6-2* (homozygous) allele whose error bar for DP3:DP4 ratio was derived from two technical replicates. Other *cslf6-2* plants were below the threshold for (1,3;1,4)-β-glucan detection therefore DP3:DP4 ratio analysis was not possible. Asterisks indicate statistically significant differences compared to the control (WT) at P < 0.01 (**) and P < 0.001 (***) using Tukey’s test in GraphPad Prism v.8.4.2. **(C)** Confocal images of transverse mature grain sections (10X) labelled with BG1 antibody in *cslf6* alleles (*cslf6-2* and *cslf6-2/+*) and cv. Golden Promise (WT). (1,3;1,4)-β-Glucan was detected in green fluorescence using Alexa Fluor 488 (BG-1). Cell wall structure was observed by calcofluor (CF) counterstaining, in blue. Below, light microscope images (5X) of the same lines. Starch content was detected by Lugol’s (LG) iodine staining. Toluidine blue O (TB) was used for differentially staining primary and secondary cell walls. **(D)** Confocal images of developing grain (15 DPA) in *cslf9* alleles (*cslf9-1*, *cslf9-2* and *cslf9-3*) and WT (transverse grain sections, 10X). Below, light microscope images (5X) of the same *cslf9* lines. (1,3;1,4)-β-Glucan and other cell wall components were detected as described for *cslf6* mutants above. Scale bars are 1mm.

### Mutations in *HvCslF6* influence grain (1,3;1,4)-β-glucan content

Grain from *cslf6-2* (homozygous) mutants almost completely lack (1,3;1,4)-β-glucan (0.11% w/w ± 0.04), whereas grain from *cslf6-2/+* (heterozygous lines) have intermediate levels of (1,3;1,4)-β-glucan (1.45% w/w ± 0.31) compared to the WT control (5.00% w/w ± 0.03) (Figure 3A). There was no significant difference between (1,3;1,4)-β-glucan content of grain from *cslf9* homozygous mutants (4.73% w/w ± 0.37, on average) compared to WT grain regardless of the mutation present (Figure 3A). We observed a consistent DP3:DP4 ratio ranging from 2.6 to 3.1 in mature grain flour samples across all lines analysed. This DP3:DP4 ratio was detected despite differences in the predicted effects of the CRISPR/Cas9 induced mutations on protein length and structure (Figure 3B). The method was sensitive enough to analyse a single *cslf6-2* (knock-out) mutant with 0.20% (w/w) ± 0.01 (1,3;1,4)-β-glucan content, showing a 3.1:1 ± 0.01 DP3:DP4 ratio. Other *cslf6-2* knock-out lines were below the threshold for (1,3;1,4)-β-glucan detection, therefore DP3:DP4 ratio analysis was not possible. Three *cslf6-2/+* (heterozygous) mutants carrying the same mutation as *cslf6-2* had an average DP3:DP4 ratio of 2.89:1 ± 0.01, which is not significantly different to the WT controls (p = 0.285, Tukey’s test).

### Distribution of (1,3;1,4)-β-glucan in mature and developing grains is comparable to wild type except in *cslf6*

To assess the distribution of (1,3;1,4)-β-glucan in WT grain compared to lines carrying induced mutations, we used immunolabelling with the BG1 antibody (Burton *et al.*, 2011). Developmental stages for analysis were selected based on the transcript abundance of the target genes; for *HvCslF6* we sampled mature grain, and for *HvCslF9* we sampled grain at 15 DPA, as this was just after the peak of expression of this gene (Burton *et al.*, 2008, Garcia-Gimenez *et al.*, 2019).

In thin sections of mature WT grain, (1,3;1,4)-β-glucan was detected in the starchy endosperm and cell walls of aleurone layers. In contrast, and as expected based on the (1,3;1,4)-β-glucan assay, labelling was completely absent in the endosperm of the *cslf6-2* (homozygous) grain. We observed weak fluorescence in the cell walls of sub-aleurone cells and outer endosperm cells (Figure 3C and 4B), but this may be partially due to autofluorescence of phenolic acids (Figure S8). *cslf6-2/+* (heterozygous) grain showed intermediate levels of (1,3;1,4)-β-glucan labelling across the aleurone and endosperm cell walls compared to WT grain (Figure 3C and 4). These results are consistent with the quantification of (1,3;1,4)-β-glucan in these lines (Figure 3A). Differences in grain shape and structure between *cslf6* mutants and WT were apparent based on calcofluor counterstaining (Figure 3C and D). Grain sections were also stained with iodine to detect starch granules revealing that both homozygous and heterozygous mutants showed more even, compact staining than WT in mature endosperm tissues. Additionally, a potential difference in polysaccharide distribution was observed in the sub-aleurone and young endosperm tissue of the *cslf6-2* (homozygous) mutant as revealed by intense staining with Toluidine blue (TB) (Figure 3C).

**Figure 4.**
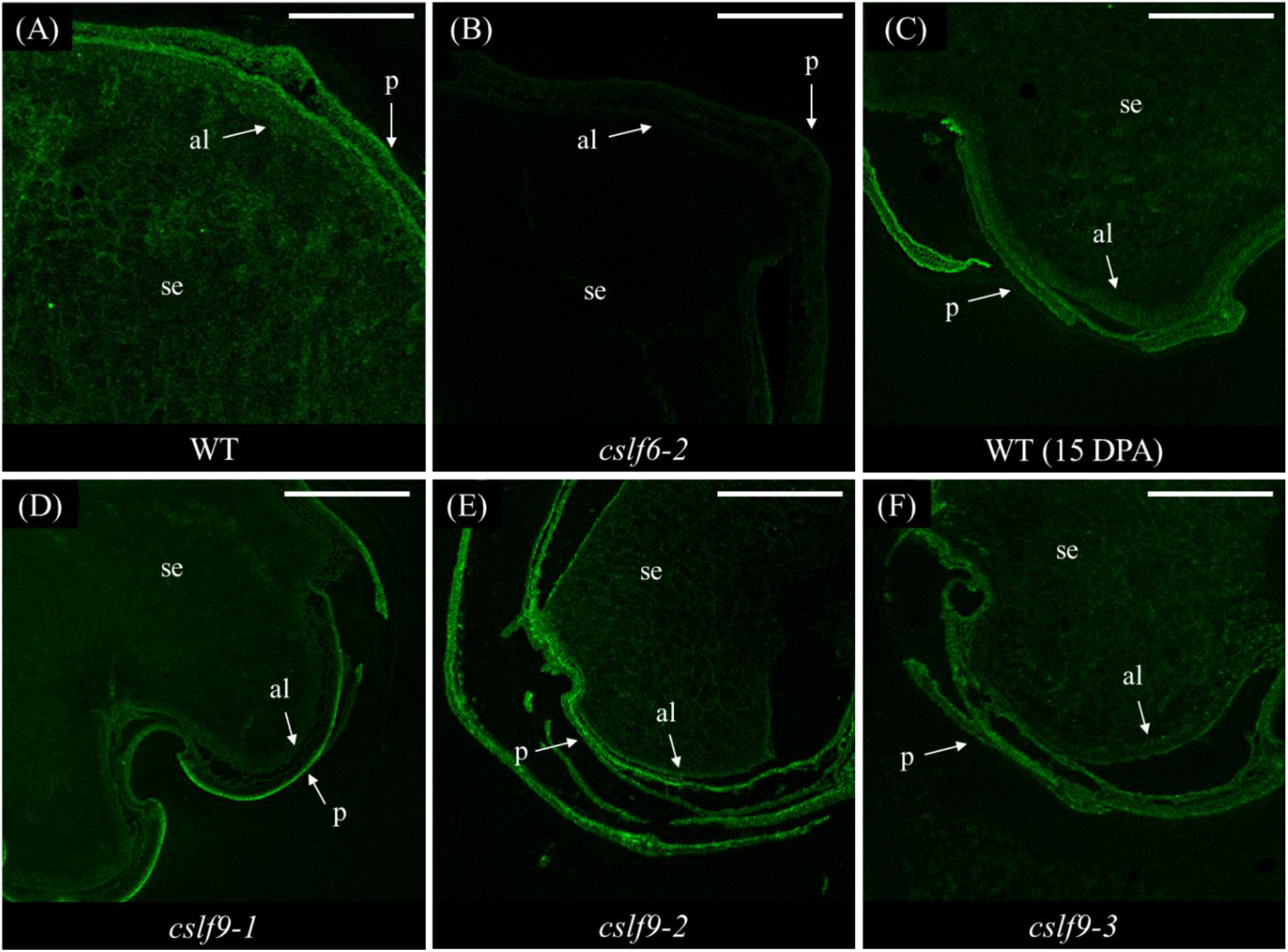
Confocal images (10X) of transverse grain sections labelled with BG1 antibody. **(A)** mature grain of cv. Golden Promise (WT), **(B)***cslf6-2* mature grain, **(C)** WT, 15 DPA, **(D)***cslf9-1*, 15 DPA, **(E)***cslf9-2*, 15 DPA **(F)***cslf9-3*, 15 DPA. (1,3;1,4)-β-Glucan was detected in green fluorescence using Alexa Fluor 488 (BG-1). The pericarp (p), aleurone (al) and starchy endosperm tissues are indicated. Scale bar = 0.5 mm.

Developing grains (15 DPA) from *cslf9* mutants (all homozygous) exhibited a similar (1,3;1,4)-β-glucan labelling pattern (i.e. distribution) compared to WT grain (Figure 3D and 4). However, this was accompanied by a reduction in calcofluor staining intensity in the *cslf9-2* mutant, the outer layers of the *cslf9-3* mutant (both of which contain a premature stop codon) and to a lesser extent in the *cslf9-1* mutant (which contains a 39 bp deletion) (Figure 3D). Further characterization of *cslf9* alleles by linkage analysis (Table S5; Method S3) showed a 24.11% ± 1.56 reduction in cellulose content compared to WT grain which was also verified by crystalline cellulose biochemical assay (18.27% ± 0.62 reduction). Moreover, *cslf9-2* and *cslf9-3* knock-out mutants appear to have a different distribution of starch and cell wall polysaccharide staining in transverse grain sections compared to WT grain. A stronger TB stain in the outer endosperm cells of mutants was observed whereas in WT grain, a darker blue colour was seen around the central grain area, indicating a potential difference in polysaccharide distribution (Figure 3D).

### Mutations in *HvCslF6* and *HvCslF9* alter grain polysaccharide composition

The differences in staining intensity indicated that polysaccharide distribution, amount and/or cell structure may be altered between *cslf6* and *cslf9* mutant grain and WT. To assess the monosaccharide composition of these mutant lines mild acid hydrolysis was used on mature grain flour. Standard conditions were followed such that only non-cellulosic polysaccharides (but not cellulose) would be hydrolysed. Consistent with previous studies, the most abundant non-cellulosic monosaccharides detected in WT non-destarched mature grain flour samples were glucose, followed by xylose, arabinose and galactose (Table 2). These represent the building blocks of the three most abundant non-cellulosic polysaccharides reported in barley grain; starch, (1,3;1,4)-β-glucan and arabinoxylan (Wilson *et al.*, 2012). Significant differences in non-cellulosic monosaccharide composition were identified between WT and mutants, particularly in glucose levels between WT (54.04 ± 0.93 % w/w) and *cslf6-2* (36.85 ± 0.89 % w/w), *cslf9-2* (43.37 ± 1.77 % w/w) and *cslf9-3* (40.18 ± 2.13 % w/w) alleles (Table 2). The reduced levels of glucose in *cslf6-2* mirrored the reduction in (1,3;1,4)-β-glucan content (Figure 3A). The reduction of glucose observed in *cslf9-2* and *cslf9-3*, both of which carry a premature stop codon, was not detected in *cslf9-1* that carries the 39 bp deletion. Moreover, no significant difference was detected in (1,3;1,4)-β-glucan content (% w/w) in any of the *cslf9* mutant lines compared to WT (Figure 3A), suggesting that the reduction in glucose may reflect a change in starch abundance (Table S5). Small but significant increases in xylose and arabinose content were identified in *cslf6-2* and *cslf9-3* lines compared to WT (Table 2). Due to the altered non-cellulosic monosaccharide composition of grain from *cslf9* alleles compared to WT, total starch content was analysed in these lines (Table S5). A significant reduction in total starch was observed across all *cslf9* alleles (50.90 ± 0.61 % w/w *cslf9-1;* 45.62 ± 0.19 % w/w *cslf9-2* and 40.24 ± 0.45 % w/w *cslf9-3*) compared to WT grain (59.32 ± 0.97 % w/w, p < 0.001, Tukey’s test; Table S5).

**Table 2.** Non-cellulosic monosaccharide mean values (% w/w ± SE) in gene-edited *cslf/h* mutants (T_2_). Asterisks indicate statistically significant differences compared to the control (WT) at p < 0.05 (*), p < 0.01 (**) and p < 0.001 (***) determined by one-way ANOVA for each monosaccharide independently (p < 0.001, glucose; p < 0.001, xylose; p < 0.001, arabinose and p < 0.01, galactose) followed by Tukey’s post-hoc test using GraphPad Prism v.8.4.2 (GraphPad Software, San Diego, California USA).

| Genotype | Glucose | Xylose | Arabinose | Galactose |
| --- | --- | --- | --- | --- |
| WT | 54.04 ± 0.93 | 4.66 ± 0.08 | 2.21 ± 0.01 | 0.25 ± 0.01 |
| *cslf3-1* | 50.98 ± 0.92 | 4.46 ± 0.06 | 2.24 ± 0.01 | 0.26 ± 0.01 |
| *cslf3-2* | 45.33 ± 0.78 * | 4.72 ± 0.04 | 2.45 ± 0.02 | 0.26 ± 0.00 |
| *cslf6-2* | 36.85 ± 0.89 *** | 5.73 ± 0.20 *** | 2.61 ± 0.06 *** | 0.35 ± 0.02 ** |
| *cslf6-2/+* | 41.28 ± 1.23 *** | 5.04 ± 0.17 | 2.51 ± 0.04 ** | 0.31 ± 0.01 |
| *cslf9-1* | 51.40 ± 1.01 | 4.74 ± 0.12 | 2.30 ± 0.02 | 0.28 ± 0.01 |
| *cslf9-2* | 43.37 ± 1.77 *** | 5.06 ± 0.09 | 2.39 ± 0.02 | 0.29 ± 0.02 |
| *cslf9-3* | 40.18 ± 2.13 *** | 6.17 ± 0.35 *** | 2.84 ± 0.13 *** | 0.29 ± 0.01 |
| *cslh1-1* | 47.36 ± 1.56 | 4.99 ± 0.03 | 2.52 ± 0.01 | 0.25 ± 0.00 |
| *cslh1-2* | 48.60 ± 1.10 * | 4.92 ± 0.10 | 2.44 ± 0.04 * | 0.30 ± 0.01 |
| *cslh1-3* | 46.46 ± 0.92 | 5.13 ± 0.02 | 2.53 ± 0.02 | 0.29 ± 0.02 |

Taken together, these findings suggest that several of the CRISPR-induced mutations in *HvCslF6* and *HvCslF9* appear to have an effect on the abundance of cellular carbon stores in the barley grain.

## Discussion

Multiple genes have been reported to play a role in (1,3;1,4)-β-glucan biosynthesis. These include genes from the *CslF* family, of which *CslF6* has a prominent role (Burton *et al.*, 2006). Gain of function assays in tobacco and *Arabidopsis* indicate that *CslF2*, *CslF4*, *CslH1* and *CslJ* may also contribute to (1,3;1,4)-β-glucan biosynthesis (reviewed in Amos & Mohnen, 2019), while QTL and GWAS studies in barley have hypothesised roles for other *CslF* family members (Han *et al.*, 1995; Houston *et al.*, 2014; Molina-Cano *et al.*, 2007). Despite this, loss-of-function mutants have only confirmed the function of *CslF6* in barley and rice (Taketa *et al.*, 2012; Cu *et al.*, 2016; Vega-Sanchez *et al.*, 2012), while functional evidence supporting a role for the other genes has been lacking. In this study a collection of mutants was successfully generated for several members of the *HvCslF*/*H* subfamilies of GT2 enzymes, and a subset of these lines were characterised at both the phenotypic and molecular level. This pool of gene-edited mutants represents a resource for the functional assessment of these and other cell wall-related genes.

To initially test our gene editing system, we targeted the *HvCslF6* gene. Taketa *et al.* (2012) previously reported *cslf6* alleles containing amino acid substitutions in barley *betaglucanless (bgl)* mutants. Here we aimed to generate a true knock-out allele by targeting the first exon of *HvCslF6*. Although relatively few lines were recovered, we successfully generated the *cslf6-2* allele that contains a premature stop codon and was therefore of interest for further characterisation. Grain (1,3;1,4)-β-glucan quantification in *cslf6-2* mutants (homozygous for a 1 bp insertion leading to a premature stop codon in the first exon) confirmed the severe reduction of this polysaccharide in mature grain (0.11% w/w ± 0.04) while intermediate (1,3;1,4)-β-glucan levels were detected in *cslf6-2/+* heterozygous lines (1.45% w/w ± 0.31). Low (1,3;1,4)-β-glucan content has traditionally been desired in malting barley breeding programs to ensure efficient filtration processes during brewing via decreased mash and wort viscosity. Although the gene-edited *cslf6* mutants reported here show extremely low levels of grain (1,3;1,4)-β-glucan, they are unsuitable for malting due to poor agronomic traits including decreased TGW, flatter and longer grains, and low germination rate. Future gene-editing efforts might either focus on targeting specific residues in the *HvCslF6* coding sequence closer to the 3’end (Dimitroff *et al.*, 2016; Jobling, 2015) or modulating transcription of the *HvCslF6* gene via regulatory regions to specifically control (1,3;1,4)-β-glucan content in the grain. Regardless, our results demonstrate the effectiveness of CRISPR/Cas9-based gene editing technology and confirm the major role of *HvCslF6* in grain (1,3;1,4)-β-glucan biosynthesis, as previously reported in barley (Burton *et al.*, 2011; Hu *et al.*, 2014; Taketa *et al.*, 2012), wheat (Nemeth *et al.*, 2010) and rice (Vega-Sanchez *et al.*, 2012).

Despite the differences in alleles generated and genetic background used, a similar phenotype to the naturally-occurring barley *betaglucanless* (*bgl*) mutant (Taketa *et al.*, 2012) was observed in our *cslf6-2* knockouts; a reduction in plant height, vigour and spike development. Such extreme phenotypes are to be expected given that *HvCslF6* is expressed in a wide range of tissues and developmental stages (Burton *et al.*, 2008). Moreover, *HvCslF6* is the main gene responsible for the synthesis of (1,3;1,4)-β-glucan, which has an integral role in the primary cell wall of barley. When DP3:DP4 ratios were quantified, no differences were observed in *cslf6* homozygous and heterozygous mutant grain compared to WT. Burton *et al.* (2011) described a reduced DP3:DP4 ratio in grain of *HvCslF6* over-expressing lines, affecting (1,3;1,4)-β-glucan solubility. Although a higher DP3:DP4 ratio was noticed in *cslf6-2/+* heterozygous mutants, this was not significantly different from controls (p = 0.285, Tukey’s test). Immunohistochemical detection of (1,3;1,4)-β-glucan confirmed the absence of this polysaccharide in mature grain of *cslf6-2* homozygous mutants whereas a weak labelling was detected across aleurone and endosperm tissues in heterozygous lines. It is known that an inverse relationship exists between (1,3;1,4)-β-glucan and starch content in grains of several grass species, including barley, based on comparative studies across cereal species with contrasting levels of this polysaccharide (Lim *et al.*, 2019; Trafford *et al.*, 2013). The precise mechanism/s that regulate this relationship remain unknown. Sections of *cslf6* mutant grain (homozygous and heterozygous) analysed for starch distribution suggest a stronger and more compact staining in sub-aleurone layers of knock-out and heterozygous lines, potentially due to lower levels of (1,3;1,4)-β-glucan.

In addition to confirming the utility of the gene editing system, the *cslf6-2* lines described here provide an important “null” background for future experiments. These lines can be used to either investigate the molecular, spatial and temporal details of *HvCslF6* or other gene function in grain (1,3;1,4)-β-glucan synthesis. Previous studies of wheat addition lines carrying *HvCslF6* revealed a 60% increase in grain (1,3;1,4)-β-glucan content, although when compared to barley controls, this only reached 20% of the (1,3;1,4)-β-glucan level in the grain (Cseh *et al.*, 2013). This indicates that other activities must be present in barley that support (1,3;1,4)-β-glucan synthesis, and these could potentially be revealed by further analysis of *cslf6-2* knock-out lines. It is known that *HvCesA1*, *HvCesA2* and *HvCesA6*, members of the *Cellulose synthase* gene subfamily within the GT_2_ Family are co-expressed with *HvCslF6* (Burton and Fincher 2009; Wilson *et al.*, 2012). It currently remains unclear whether their function or regulation is compromised through lack of functional HvCSLF6 in *cslf6-2* lines. The *cslf6-2* knock-out alleles also provide an opportunity to create double mutants, for example with *cslh1* lines carrying frameshift mutations, to assess the contribution of multiple members of the *Csl* gene family to cell wall composition in leaf, grain and other tissues.

Unlike *HvCslF6*, previous studies have not reported loss-of-function phenotypes for the *HvCslF9* gene. *HvCslF9* expression in developing grain is generally low but shows a prominent peak at around 8 DPA (Burton *et al.*, 2008). This peak coincides with initiation of (1,3;1,4)-β-glucan synthesis, which by 10 DPA is uniformly distributed throughout the cellularised endosperm (Wilson *et al.*, 2006). One possibility is that *HvCslF9* contributes to (1,3;1,4)-β-glucan synthesis in a temporally and/or spatially restricted manner. Our data confirm that *HvCslF9* has a role in barley grain development. Knockout alleles exhibited pleiotropic changes in grain morphology, monosaccharide, polysaccharide and starch composition, but only minor changes in grain (1,3;1,4)-β-glucan content. This was confirmed in four independent knockout mutants, two lines carrying a 5 bp deletion (*cslf9-2*) and another two carrying a 1 bp insertion (*cslf9-3*), all leading to premature stop codons. Similar results were obtained for *cslf9-1* mutants carrying a 39 bp deletion in the first exon.

Immunolabelling was used to assess whether a loss of *HvCslF9* leads to temporal and/or spatial differences in (1,3;1,4)-β-glucan deposition during earlier stages of grain development. BG-1 labelling at 15 DPA confirmed the presence of (1,3;1,4)-β-glucan in the endosperm and aleurone layers of *cslf9* knock-out lines with similar distribution compared the WT. The detection of (1,3;1,4)-β-glucan in mid and mature grain samples suggests that *HvCslF9* might not contribute to mature grain (1,3;1,4)-β-glucan content. However, as we did not characterise (1,3;1,4)-β-glucan chain length or fine structure we cannot rule out a role for *HvCslF9* in these aspects of this polysaccharide. Various QTL and GWAS reports (Han *et al.*, 1995; Molina-Cano *et al.*, 2007; Houston *et al.*, 2014) identified a region on chromosome 1H involved in (1,3;1,4)-β-glucan content, which encompasses *HvCslF9* and a (1,3;1,4)-β-glucan endohydrolase (*HvGlbI*) involved in the hydrolysis of this polysaccharide. This QTL could also have an impact on malt (1,3;1,4)-β-glucan modification (Stuart *et al.*, 1988; Slakeski & Fincher 1992) and has been reported in QTL studies that quantified malt (1,3;1,4)-β-glucan content (Han *et al.*, 1997, 1995; Szűcs *et al.*, 2009; Ullrich *et al.*, 1997). Evidence presented in the current study suggests that *HvCslF9* is unlikely to influence mature grain (1,3;1,4)-β-glucan content in the Golden Promise cultivar, and hence, *HvCslF9* may not be the gene underlying this 1H QTL. Alternatively, *HvCslF9* may contribute to (1,3;1,4)-β-glucan biosynthesis only in selected genotypes, at an earlier stage of grain development and/or in a defined grain tissue compartment not investigated here.

Despite there being no significant difference in the amount of grain (1,3;1,4)-β-glucan present compared to WT, the *cslf9* mutants exhibited a lower TGW compared to WT grain. In the case of the two lines containing premature stop codons in *HvCslF9* this reduction in TGW was coupled with a decrease in total starch, cellulose and arabinoxylan compared to WT. Subtle spatial differences in staining intensity were identified in the same lines using iodine (starch), toluidine blue (general cell walls) and calcofluor white (general cell walls including cellulose, callose, and (1,3;1,4)-β-glucan) stains. These changes suggest that *HvCslF9* may contribute to polysaccharide synthesis in specific regions of the grain, and or specific time points during development, and this warrants further investigation via molecular and biochemical assays. A recent publication using transient heterologous expression in *Nicotiana benthamiana* leaves indicated that the *HvCslF3* and *HvCslF10* genes may be involved in the synthesis of a novel linear (1,4)-β-xyloglucan that consists of (1,4)-β-linked glucose and xylose residues (Little *et al.*, 2019). The same study investigated activity of *HvCslF9* but failed to identify any clear biosynthetic activity in the heterologous system. Combined with the results described by Little *et al.* (2018, 2019), our data agree with the hypothesis that some members of the *CslF/H* gene families could either influence or be involved in the synthesis of polysaccharides other than (1,3;1,4)-β-glucan.

The final two genes examined were *HvCslF3* and *HvCslH1*, which are expressed at very low levels during grain development (Burton *et al.*, 2008, Doblin *et al.*, 2009). Therefore, as expected, frameshift mutations affecting *HvCslF3* and *HvCslH1* did not significantly alter either grain monosaccharide abundance or (1,3;1,4)-β-glucan content (Table 3, S4). Highest mRNA abundance is found for *HvCslF3* in roots (Aditya *et al.*, 2015) and for *HvCslH1* in leaves (Doblin *et al.*, 2009). However, while a clear role for *CslH1* in (1,3;1,4)-β-glucan synthesis was demonstrated by Doblin *et al.* (2009) using heterologous expression in tobacco leaves, when Burton *et al.* (2011) overexpressed *HvCslF3* in barley grain there was no detectable change in (1,3;1,4)-β-glucan content. Loss-of-function mutations in either gene impacted mature grain morphology, suggesting they are likely to have a function during plant development, but changes in mature grain composition were relatively small and statistically insignificant. Further phenotypic characterisation of additional plant tissues and developmental stages will be required to determine the precise contribution of these genes to cell wall composition; whether they have an organ- or tissue-specific role in (1,3;1,4)-β-glucan biosynthesis, are impacted by compensatory mechanisms (Pérez *et al.*, 2019) or synthesise other polysaccharides such as glucoxylan (Little *et al.*, 2019).

The efficiency of CRISPR/Cas9-induced mutations in the primary transformants ranged from 5–35% in this study. Previous studies in barley reported CRISPR/Cas9-induced mutation frequencies of 10–23% (Lawrenson *et al.*, 2015), 44% (Holme *et al.*, 2017) and ~80% (Gasparis *et al.*, 2018; Kapusi *et al.*, 2017) of plants after *Agrobacterium-*mediated transformation. In other cereals such as wheat, mutant production efficiency was reported from 0.3–5% using biolistic bombardment of CRISPR/Cas9 components (Zhang *et al.*, 2016; Gil-Humanes *et al.*, 2017; Liang *et al.*, 2017; Sánchez-León *et al.*, 2018) and *Agrobacterium-*mediated transformation (Okada *et al.*, 2019). In the T_0_ plants described here, all mutants were heterozygous (monoallelic) across the targeted genes, with 83% of the lines carrying short InDels 3–4 bp upstream the PAM sequence. This location of the Cas9-induced mutation relative to the PAM sequence agrees with Cas9 cleavage pattern and error-prone NHEJ. We did not detect any homozygous mutations in primary transformants (T_0_) unlike other studies in rice (Zhang *et al.*, 2014; Zhou *et al.*, 2014).

Taken together, our collection of gene-edited alleles for *HvCslF*/*H* with knock-out and frameshift mutations represent a valuable genetic resource for studying barley cell walls and demonstrate the effectiveness and potential of CRISPR/Cas9 genome editing technology for the analysis of gene function. Importantly, our results show that members of the *CslF/H* family other than *HvCslF6* have an *in-planta* function in growth and development, providing greater opportunities to understand and optimise cell wall composition for specific downstream applications.

## Supplementary data

**Table S1.** Summary of transformation results (T_0_) across *HvCslF*/*H* genes.

**Table S2.** Allelic state of CRISPR/Cas9-induced mutations and Cas9 presence in T_1_ plants.

**Table S3.** BLAST score results for *HvCslF3*, and *HvCslH1* sgRNAs.

**Table S4.** Subset of *cslf3* and c*slh1* mutants used for grain (1,3;1,4)-β-glucan quantification.

**Table S5.** Grain composition of *cslf9* alleles (T_2_) compared to WT.

**Table S6.** sgRNA and PAM sequences for *HvCslF3, HvCslF6, HvCslF9* and *HvCslH1*.

**Table S7.** Phire and Hot Start Taq PCR conditions.

**Table S8.** Primer sequences used in this study.

**Figure S1.** Frequency of mutations in T_0_, T_1_ and T_2_ generations across the target genes.

**Figure S2.** Determination of *Cas9* copy number by hygromycin segregation test.

**Figure S3.** Images of whole plant phenotype in the *cslf6-2* homozygous mutant.

**Figure S4.** (A) Plant phenotype of *cslf3* mutants. (B) Assessment of grain characteristics.

**Figure S5.** (A) Plant phenotype of *cslh1* mutants. (B) Assessment of grain characteristics.

**Figure S6.** Representation of sgRNA + PAM sequences in the context of BLAST hits.

**Figure S7.** Vector map of pBracT_2_14m-HvCas9-HSPT-sgRNA (destination vector).

**Figure S8.** Negative controls used for grain immunolabeling of (1,3;1,4)-β-glucan.

**Method S1.** sgRNA construct assembly.

**Method S2.** Tissue fixation, embedding and immunocytochemistry

**Method S3.** Starch and cellulose quantification

## Author Contributions

KH, MRT, RAB and RW conceived this work. KH designed and generated the sgRNA constructs. JS carried out barley transformations. Abdellah Barakate adapted the genotyping method for mutation detection. GG-G and KH performed the genotypic characterisation of the CRISPR lines. GG-G carried out the grain phenotyping, (1,3;1,4)-β-glucan quantification, immunolabelling experiments and data analyses. PS processed the CRISPR lines/samples for subsequent assays. SFK and RAB carried out the grain monosaccharide and starch analyses. MSD and PH performed the linkage and cellulose analyses. The manuscript was drafted by GG-G, KH and MRT and reviewed by Abdellah Barakate, Antony Bacic, RAB, MSD, RW and GBF. All the authors read and approved the manuscript.

## Acknowledgements

We thank Diane Davidson from the Functional Genomics (FUNGEN) facility, The James Hutton Institute, UK for barley transformation. We gratefully acknowledge Dr. Alison Roberts and Dr. Kathryn Wright, The James Hutton Institute (UK), for technical assistance during immunolabelling experiments and confocal imaging. We thank Edwin R. Lampugnani for discussions. RW and Abdellah Barakate acknowledge support from ERC project 669182 ‘SHUFFLE’ to RW. KH, PS, JS and RW acknowledge support from BBSRC project BB/J014869/1 the Rural & Environment Science & Analytical Services Division of the Scottish Government. RAB, GBF, MRT, MSD and Antony Bacic acknowledge the support of the Australian Research Council Centre of Excellence in Plant Cell Walls Grant CE1101007 and MSD, PH and Antony Bacic acknowledge the support of a startup grant from La Trobe University.

