## Supplementary material for "Targeted mutation of barley (1,3;1,4)-β-glucan synthases reveals complex relationships between the storage and cell wall polysaccharide content": Figure S1

### Slide 1
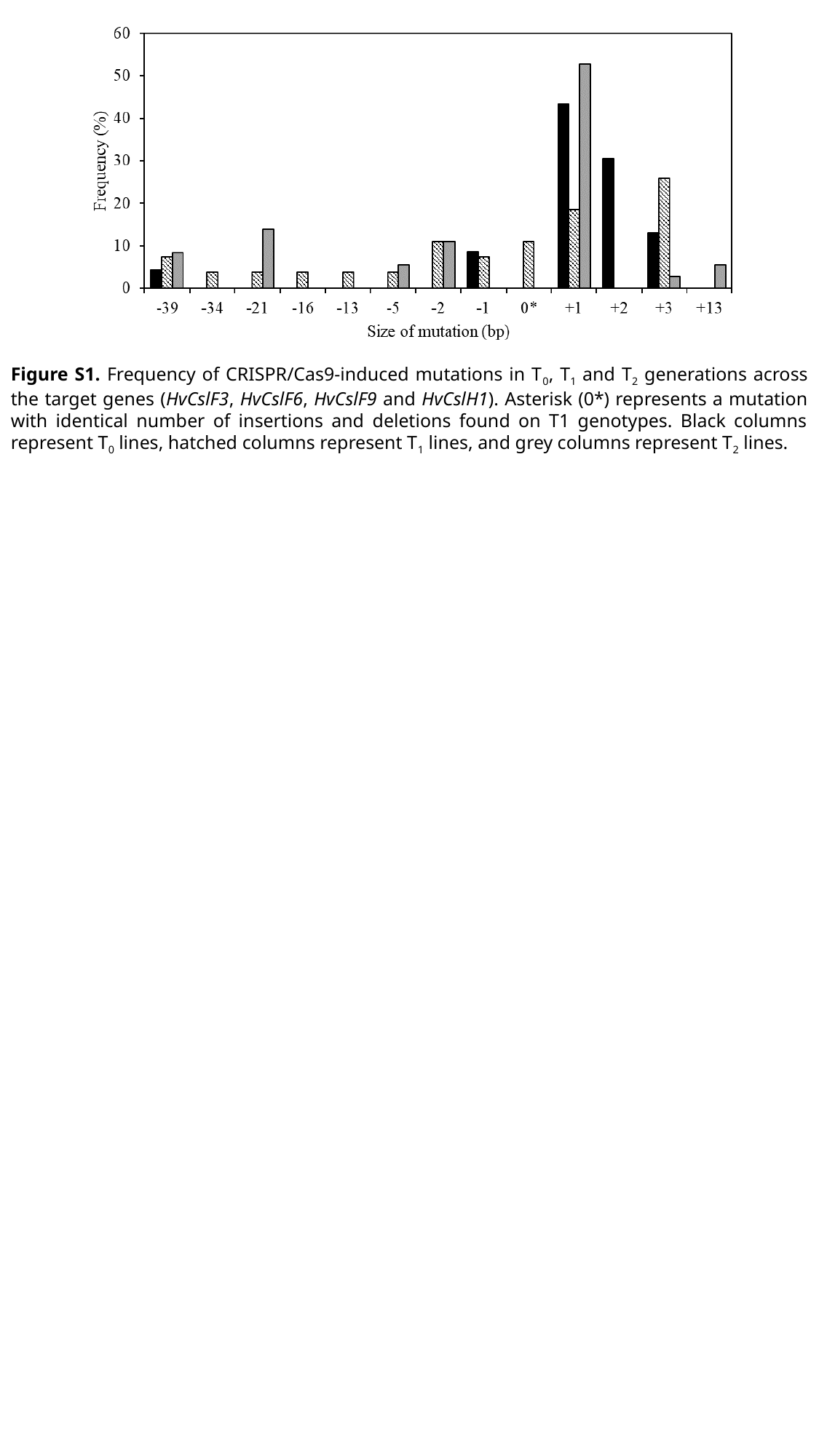

Figure S1. Frequency of CRISPR/Cas9-induced mutations in T_0_, T_1_ and T_2_ generations across the target genes (HvCslF3, HvCslF6, HvCslF9 and HvCslH1). Asterisk (0*) represents a mutation with identical number of insertions and deletions found on T1 genotypes. Black columns represent T_0_ lines, hatched columns represent T_1_ lines, and grey columns represent T_2_ lines.
