## Supplementary material for "Targeted mutation of barley (1,3;1,4)-β-glucan synthases reveals complex relationships between the storage and cell wall polysaccharide content": Figure S2

### Slide 1
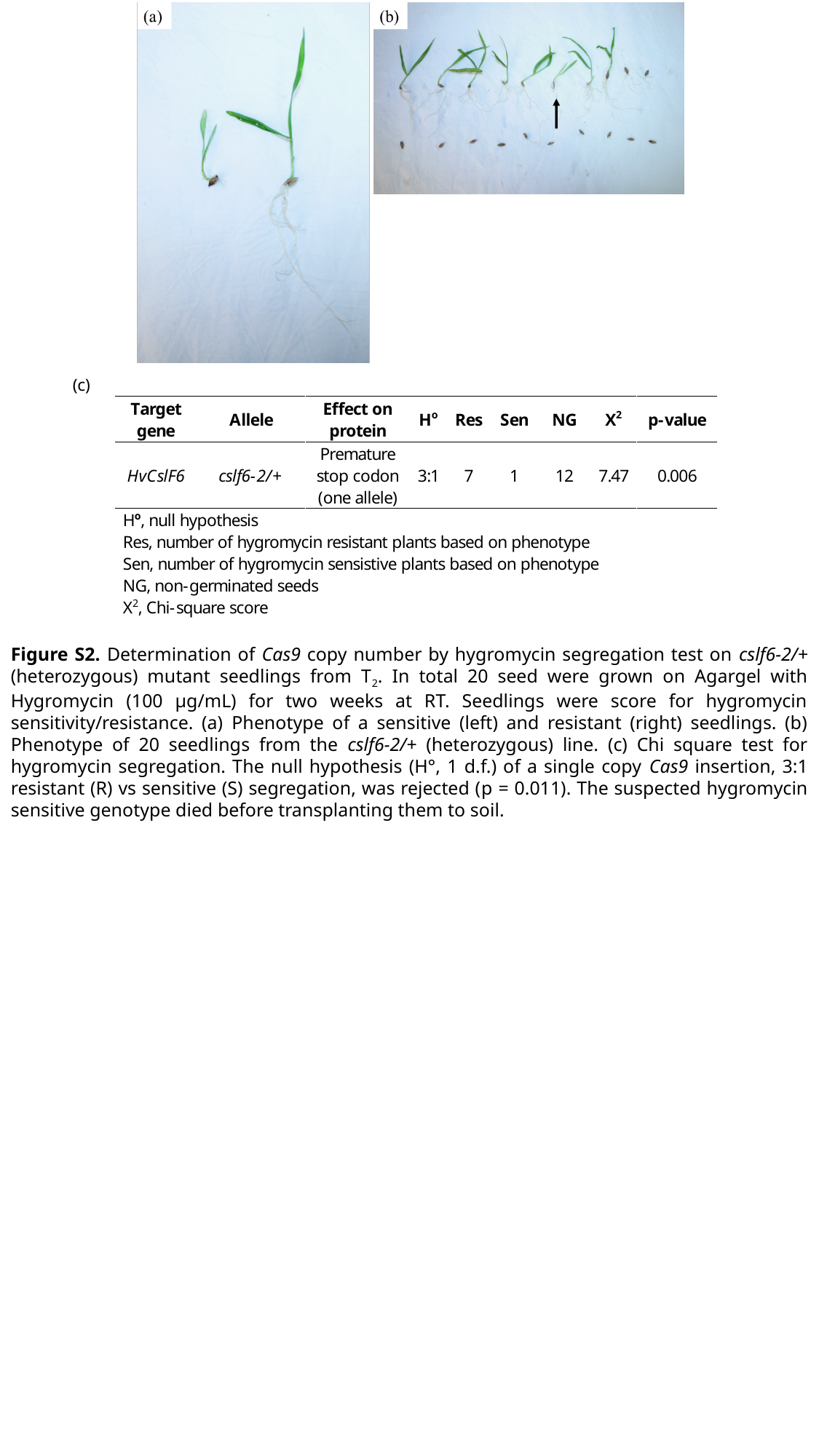

Figure S2. Determination of Cas9 copy number by hygromycin segregation test on cslf6-2/+ (heterozygous) mutant seedlings from T_2_. In total 20 seed were grown on Agargel with Hygromycin (100 µg/mL) for two weeks at RT. Seedlings were score for hygromycin sensitivity/resistance. (a) Phenotype of a sensitive (left) and resistant (right) seedlings. (b) Phenotype of 20 seedlings from the cslf6-2/+ (heterozygous) line. (c) Chi square test for hygromycin segregation. The null hypothesis (H°, 1 d.f.) of a single copy Cas9 insertion, 3:1 resistant (R) vs sensitive (S) segregation, was rejected (p = 0.011). The suspected hygromycin sensitive genotype died before transplanting them to soil.
