## Supplementary material for "Targeted mutation of barley (1,3;1,4)-β-glucan synthases reveals complex relationships between the storage and cell wall polysaccharide content": Figure S3

### Slide 1
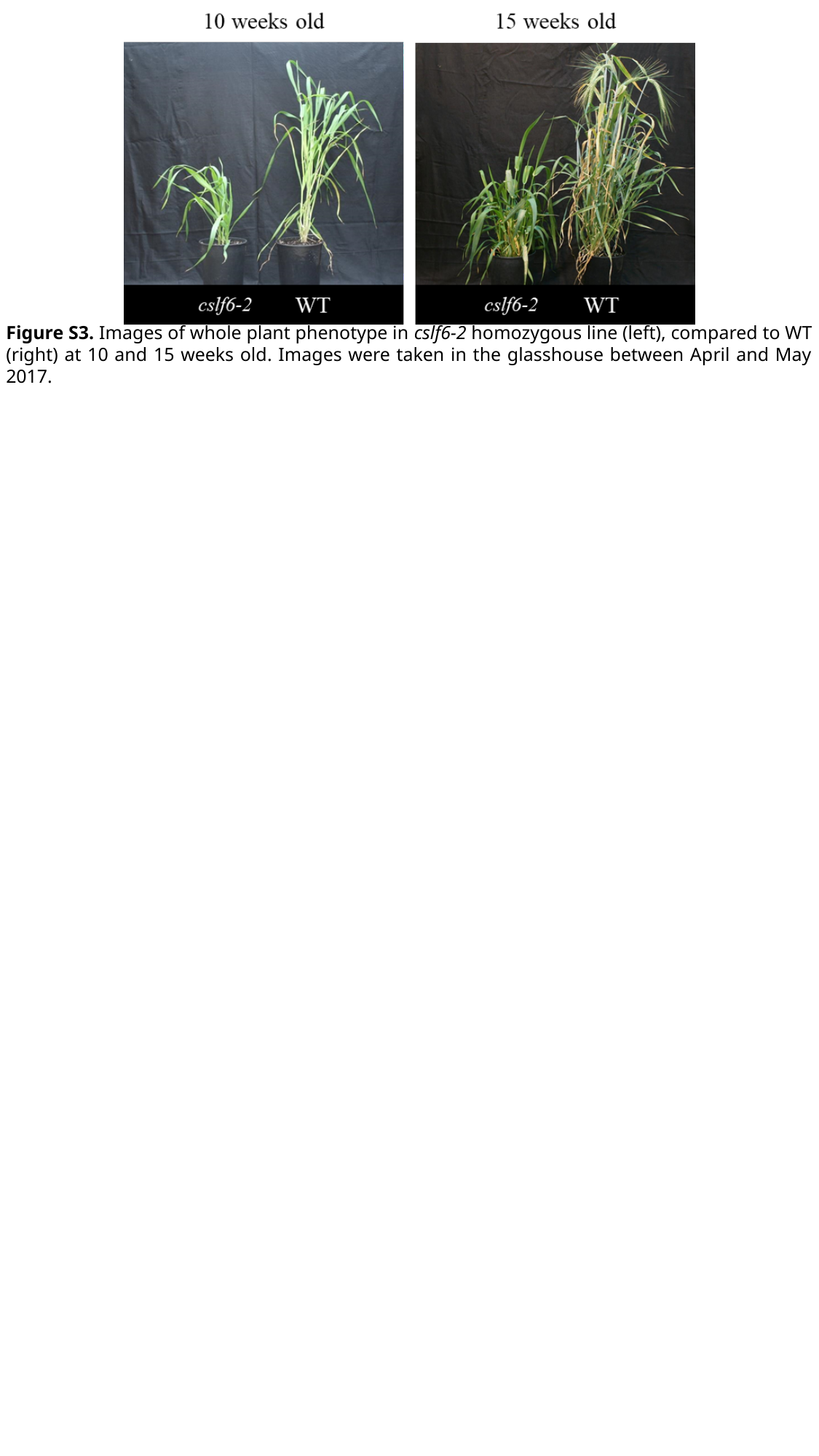

Figure S3. Images of whole plant phenotype in cslf6-2 homozygous line (left), compared to WT (right) at 10 and 15 weeks old. Images were taken in the glasshouse between April and May 2017.
