## Supplementary material for "Targeted mutation of barley (1,3;1,4)-β-glucan synthases reveals complex relationships between the storage and cell wall polysaccharide content": Figure S4

### Slide 1
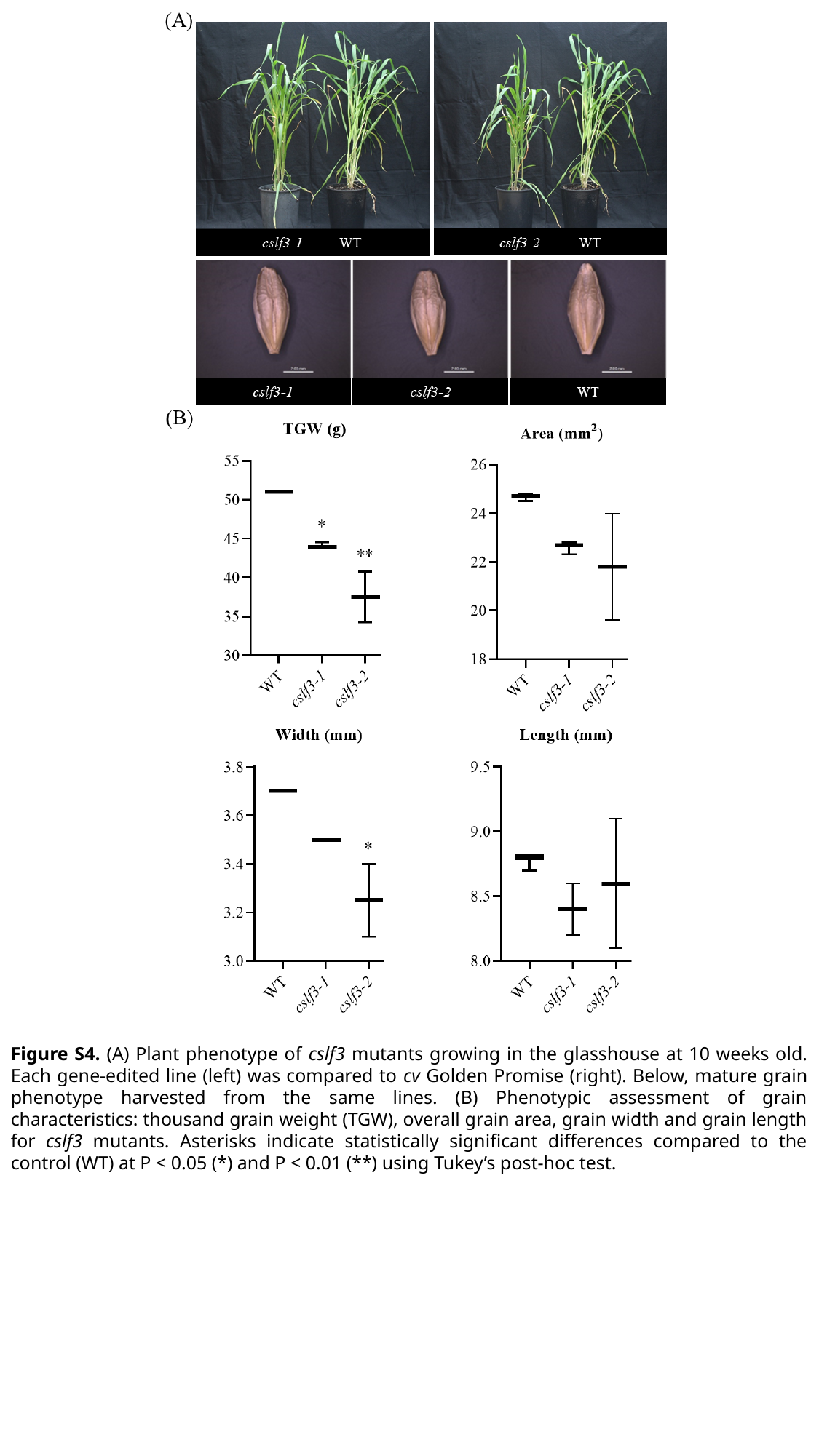

Figure S4. (A) Plant phenotype of cslf3 mutants growing in the glasshouse at 10 weeks old. Each gene-edited line (left) was compared to cv Golden Promise (right). Below, mature grain phenotype harvested from the same lines. (B) Phenotypic assessment of grain characteristics: thousand grain weight (TGW), overall grain area, grain width and grain length for cslf3 mutants. Asterisks indicate statistically signiﬁcant differences compared to the control (WT) at P < 0.05 (*) and P < 0.01 (**) using Tukey’s post-hoc test.
