## Supplementary material for "Targeted mutation of barley (1,3;1,4)-β-glucan synthases reveals complex relationships between the storage and cell wall polysaccharide content": Figure S6

### Slide 1
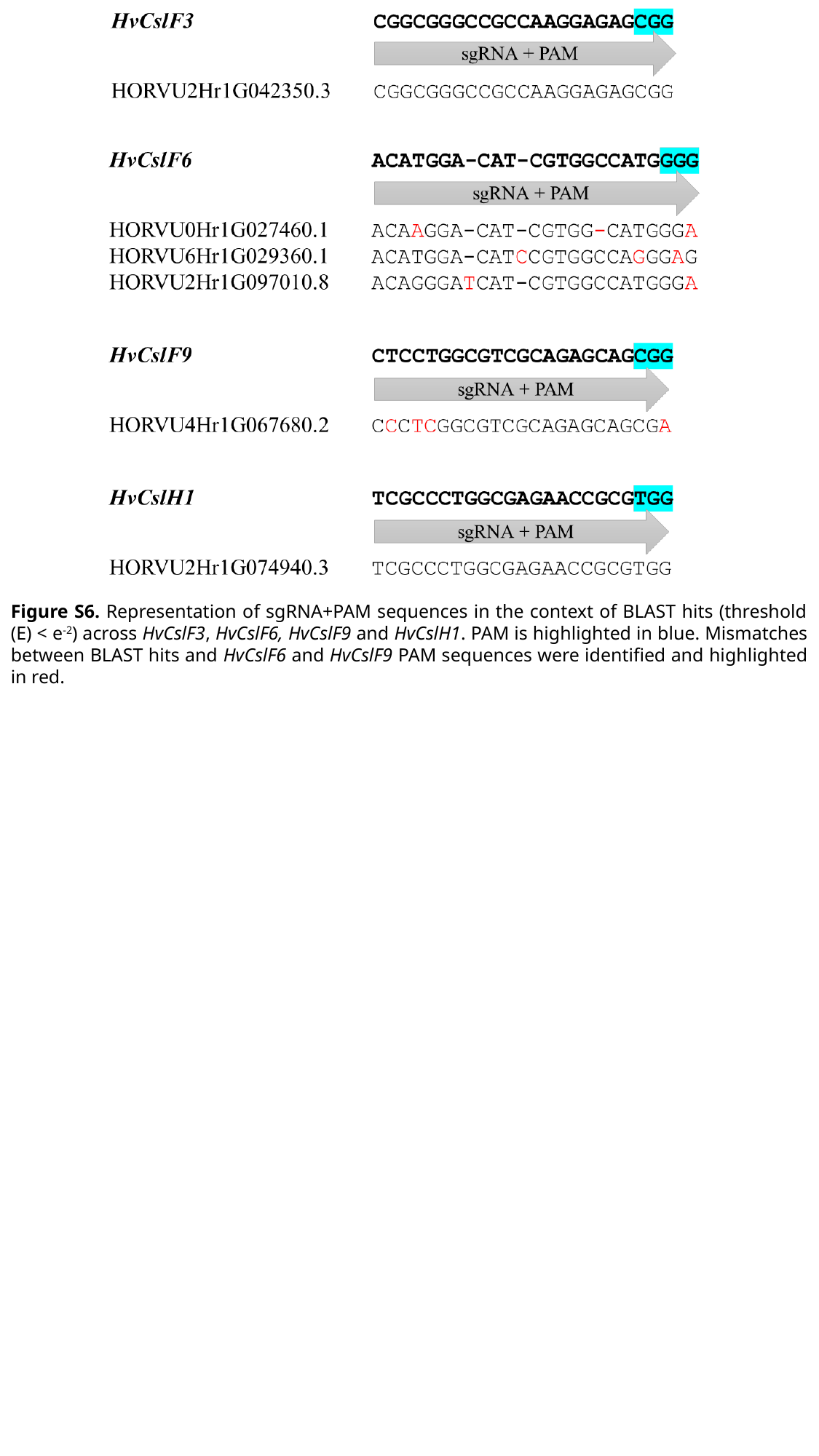

Figure S6. Representation of sgRNA+PAM sequences in the context of BLAST hits (threshold (E) < e^-2^) across HvCslF3, HvCslF6, HvCslF9 and HvCslH1. PAM is highlighted in blue. Mismatches between BLAST hits and HvCslF6 and HvCslF9 PAM sequences were identified and highlighted in red.
