## Supplementary figures and images for "Targeted mutation of barley (1,3;1,4)-β-glucan synthases reveals complex relationships between the storage and cell wall polysaccharide content"

### Figure S7

#### Slide 1
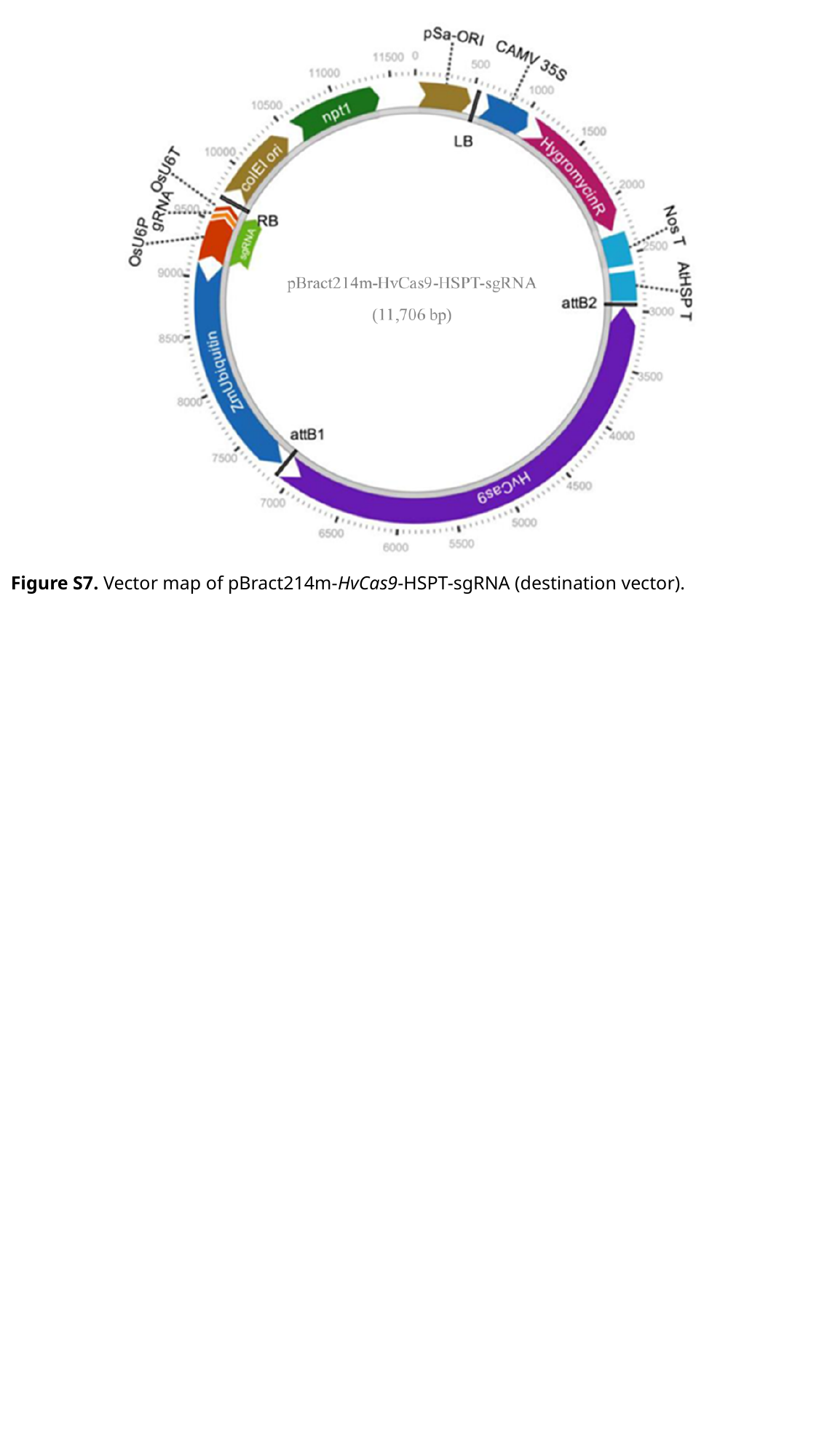

Figure S7. Vector map of pBract214m-HvCas9-HSPT-sgRNA (destination vector).
