## Supplementary material for "Targeted mutation of barley (1,3;1,4)-β-glucan synthases reveals complex relationships between the storage and cell wall polysaccharide content": Figure S8

### Slide 1
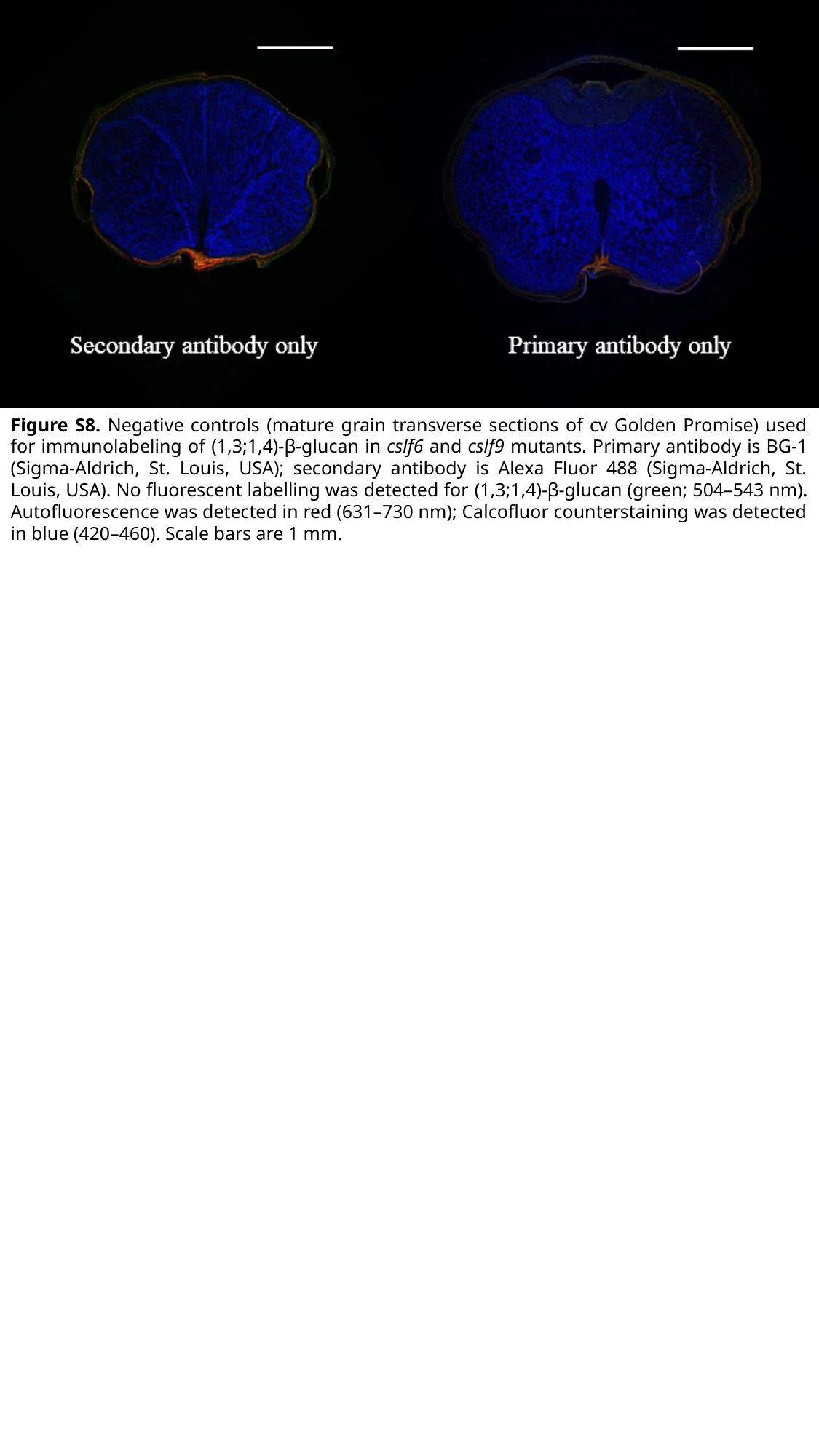

Figure S8. Negative controls (mature grain transverse sections of cv Golden Promise) used for immunolabeling of (1,3;1,4)-β-glucan in cslf6 and cslf9 mutants. Primary antibody is BG-1 (Sigma-Aldrich, St. Louis, USA); secondary antibody is Alexa Fluor 488 (Sigma-Aldrich, St. Louis, USA). No fluorescent labelling was detected for (1,3;1,4)-β-glucan (green; 504–543 nm). Autofluorescence was detected in red (631–730 nm); Calcofluor counterstaining was detected in blue (420–460). Scale bars are 1 mm.
