## Supplementary material for "Targeted mutation of barley (1,3;1,4)-β-glucan synthases reveals complex relationships between the storage and cell wall polysaccharide content": Method S2

**Method S2. Tissue fixation, embedding and immunocytochemistry**

Grain samples were fixed in 500 mL of 4% paraformaldehyde in PEM buffer solution [PEM Buffer = 0.1 M PIPES (pH 6.95), 1 mM EGTA, 1 mM MgSO_4_] for 24 hrs at 4°C. After fixation, samples were washed on a shaking platform for 30 min (3x) in sdH_2_O, (1x) 40% ethanol and (1x) 70% ethanol. Samples were transferred to a Leica TP1020 tissue processor tissue processor and waxed in Paramat Extra, in pastille form (VWR International, Lutterworth, UK). Wax blocks were transverse sectioned in a RM2265 Rotary Microtome (Leica Biosystems, Vista, USA) at 10 µm and mounted on Polysine^®^ slides (Thermo Fisher Scientific, Waltham, USA). Sections were allowed to dry for at least 12 hrs on a drying rack at 40°C to ensure well-adhered grain samples. Prior to antibody labelling, slides were de-waxed on a Tissue-Tek II^®^ Slide Staining Set (Sakura Finetek, USA) in the following 250 mL solutions: Histo-Clear II™ (2x, 30 min), 100% ethanol (2 min), 100% ethanol (5 min), 95% ethanol (2 min), 85% ethanol (2 min), 70% ethanol (2 min), 40% ethanol (2 min), sdH_2_O (2 min) and sdH_2_O (5 min). Once de-waxed, slides were returned to the drying rack for at least another 2 hrs at 40°C.

For antibody labelling of (1,3;1,4)-β-glucan slides were submerged in 100 mL of 1x PBS (phosphate-buffered saline) for 10 min using a glass staining chamber. This step was repeated using fresh 1x PBS. After, slides were transferred to 0.05 M glycine for 20 min to inactivate residual aldehyde groups. Anti-mouse primary antibody, BG1, (Sigma-Aldrich, St. Louis, USA) was diluted 1:50 in incubation buffer [1% BSA (Sigma-Aldrich, St. Louis, USA) in 1x PBS] making up 2 mL. Buffer was drained from each slide, avoiding sample drying out. 200 µL of primary antibody was applied per slide and incubated in a humidity chamber overnight at 4°C. The following day, slides were washed in incubation buffer in a glass staining chamber for 10 min. The wash step was repeated with fresh incubation buffer. Slides were transferred back to a humidity chamber and the secondary antibody was applied in dark conditions. Anti-goat secondary antibody conjugated to Alexa Fluor 488 (Sigma-Aldrich, St. Louis, USA) was diluted 1:100 in incubation buffer and 200 µL were applied per slide. Samples were incubated in the dark for 2 hrs at RT. Each batch of immunolabelling assays contained two *cv* Golden Promise negative controls (no primary antibody with secondary and vice versa) performed in parallel with the rest of the samples. After incubation, slides were washed thoroughly with 1x PBS using a Pasteur pipette and submerged in 1x PBS for 5 min. After, samples were counterstained with 0.1% calcofluor white (Sigma-Aldrich, St. Louis, USA) for 1–2 min and rinsed in sdH_2_O. For sample imaging, two to four drops of mountant [90% glycerol (Sigma-Aldrich, St. Louis, USA)] were applied and slides sealed with a glass coverslip. Samples were store in a humidity chamber at 4°C and analysed immediately after.

Barley grain sections were stained with Lugol's iodine solution (Sigma-Aldrich, St. Louis, USA) diluted 1:5 in sdH_2_O to detect starch granules. Slides were covered in Lugol's working solution (200 µL per slide) and incubated for 1 min at RT. Samples were washed with abundant sdH_2_O for 1 min and placed in the drying rack for 2 hrs at 40°C. For lignin and differential polysaccharide staining of grain sections, a 0.02 % toluidine blue O (Sigma-Aldrich, St. Louis, USA) solution was applied (200 µL per slide) and incubated for 1 min at RT. After, samples were washed repeatedly with sdH_2_O for 1 min and placed in the drying rack for at least 2 hrs at 40°C. Samples were stored in the dark at 4°C.
