## Supplementary material for "Targeted mutation of barley (1,3;1,4)-β-glucan synthases reveals complex relationships between the storage and cell wall polysaccharide content": Method S1

**Method S1. sgRNA construct assembly.**

sgRNAs for *HvCslF3*, *HvCslF6*, *HvCslF9* and *HvCslH1* were cloned into pC95-gRNA entry vector downstream of the rice U6 snRNA promoter (OsU6p) by Gibson Assembly^®^. Firstly, two 60 bases long primers (Table S3) were designed as forward and reverse oligos that are complementary to each other over the 20 bases at their 3’-ends representing the target gene specific spacer. The remaining extensions were made of 20 bases in their 5’-end overlapping the sequences flanking the insertion site in the entry vector and 20 bases at 3’-ends that restore full OsU6p-sgRNA cassette. Spacers where the first nucleotide is G were selected for correct transcriptional start from OsU6 promoter. If this was not the case, primer sequences were modified to contain an additional G as the first sgRNA nucleotide. Double strand DNA was generated by combining 5 µL of each primer (100 µM), mixed with 3.8 µL dH2O and incubated at 95°C for 5 min. After cooling to room temperature, 2 µL dNTP (2mM), 4 µL Phusion^®^ Buffer and 0.2 µL Phusion^®^ (Thermo Fisher Scientific, Waltham, USA) were added and the mix incubated at 72°C for 30 min. pC95-gRNA vector was linearized using the restriction enzyme AflII (New England Biolabs, Ipswich, USA) in the following digestion reaction: 0.25 µL AflII (20,000 units/mL), 2 µL NEBuffer 2, 13.75 µL dH2O, 4 µL pC95-gRNA (200 ng/µL) for 1h at 37°C. The digestion mix was run on a 1% agarose gel with 1 µL/ml SYBR^®^ Safe (10,000X in DMSO; Thermo Fisher Scientific, Waltham, USA), the vector band was excised according to size and purified using the QIAquick Gel Extraction Kit (Qiagen GmbH, Hilden, Germany). Following this, a Gibson Assembly^®^ reaction was performed for each sgRNA containing: 5 µL of linearized pC95-gRNA (46 ng/µL), 2 µL gRNA duplex product, 10 µL Gibson Assembly^®^ buffer-mix and 3 µL dH2O were mixed and incubated at 50°C for 60 min. The cloning mix was transformed into One Shot^™^ TOP10 chemically competent *E. coli* for multiplication and bacteria were grown overnight on ampicillin (100 µg/mL) plates. Plasmids were purified using QIAprep Spin Miniprep Kit (Qiagen GmbH, Hilden, Germany) and Sanger sequenced in-house using T7_F primer to confirm the ligation and correct sgRNA orientation. The destination vector pBract214m-HSPT-HvCas9 was linearized using StuI (New England Biolabs, Ipswich, USA) in a digestion reaction containing: 0.25 µL StuI (10,000 units/mL), 3 µL pBract214m-HSPT-HvCas9 (140 ng/µL), 2µL NEBuffer 2, 14.75 µL dH2O for 1 h at 37°C. pBract214m-HSPT-HvCas9 was dephosphorylated by adding 1 µL of Antarctic Phosphatase (5000 units/mL; New England Biolabs, Ipswich, USA) and 2 µL Antarctic Phosphatase Reaction buffer (1X) to the restriction digest and incubated at 37°C for 30 min.

sgRNAs were excised from pC95-gRNA by restriction enzyme digest with EcoRV (New England Biolabs, Ipswich, USA), in the following reaction: 0.25 µL EcoRV (20,000 units/mL), 2 µL NEBuffer n°3, 5 µL pC95-gRNA (10 ng/µL), 12.75 µL dH2O. The reaction was incubated at 37°C for 1 hour. Digestion reactions for both, sgRNA and destination vector pBract214m-HSPT-HvCas9 were run on a 1% agarose gel with 1 µL/ml SYBR^®^ Safe (10,000X in DMSO) and the appropriate bands identified according to size, excised and purified as described above. Concentration of recovered DNA were measured using NanoDrop 2000 (Thermo Fisher Scientific, Waltham, USA) and fragments were then ligated using T4 DNA ligase (Promega, Madison, USA). T4 ligation reaction contained: 3 µL pBract214m-HSPT-HvCas9 (12 ng/µL), 12 µL gRNA (15 ng/µL), 2 µL DNA Ligase Reaction Buffer (1X), 1µL T4 DNA Ligase (400,000 units/mL). The reaction was incubated at RT overnight, followed by a heat inactivation at 65°C for 10 min. After, *E. coli* was transformed with the resulting construct. Plasmids were purified and confirmed by Sanger sequencing (as previously described) using OsU6p specific forward primer.
