## Supplementary material for "Targeted mutation of barley (1,3;1,4)-β-glucan synthases reveals complex relationships between the storage and cell wall polysaccharide content": Method S3

**Method S3. Starch and cellulose quantification**

Total starch content was measured on 40 mg of flour, using a small-scale version of the Megazyme Total Starch Assay (K-TSTA) kit (McCleary et al., 1994) with quadruplicate technical replication. Cellulose content was quantified by two independent methods, monosaccharide linkage analysis and acetic/nitric acid-resistant crystalline cellulose based on Updegraff (1969) method based on single biological replicates per *cslf9* mutation. For the linkage analysis, milled samples (300 mg) were first treated with porcine α-amylase then washed extensively with 1% w/v sodium dodecyl sulphate (SDS) in 70% (v/v) ethanol to remove any remaining starch and protein, as verified by iodine/KI staining (Pettolino *et al*. 2012). Methylation analysis of the de-starched alcohol insoluble residue (AIR) samples was conducted according to the method outlined in Pettolino *et al*. (2012). Briefly, duplicate samples (10 mg) were subjected to two rounds of methylation prior to trifluoroacetic acid (TFA) hydrolysis followed by reduction and acetylation to generate partially methylated alditol acetates (PMAAs). These were then quantified by GC-MS and cellulose composition deduced as described in Pettolino *et al.* (2012). Crystalline cellulose content (acetic/nitric acid-resistant) was verified using the method of Updegraff (1969) using two technical replicates.
